# WNT vampirization by glioblastoma leads to tumor growth and neurodegeneration

**DOI:** 10.1101/428953

**Authors:** Marta Portela, Varun Venkataramani, Natasha Fahey-Lozano, Esther Seco, Maria Losada-Perez, Frank Winkler, Sergio Casas-Tintó

## Abstract

Glioblastoma (GB) is the most lethal brain tumor due to its high proliferation, aggressiveness, infiltration capacity and resilience to current treatments. Activation of the Wingless-related-integration-site (WNT) pathway is associated with a bad prognosis. Using *Drosophila* and primary xenograft models of human GB, we describe a mechanism that leads to the activation of WNT signaling [Wingless (Wg) in *Drosophila*] in tumor cells. GB cells display a network of tumor microtubes (TMs) which enwraps neurons, accumulates Wg receptor Frizzled1 (Fz1), and, thereby, actively depletes Wg from the neurons. Consequently, GB cells proliferate due to β-catenin activation, and neurons degenerate due to Wg signaling extinction. This novel view explains both neuron-dependent tumor progression, and also the neural decay associated with GB.

## Introduction

The evolution of glioblastoma (referred here as GB) is accompanied by broad neurological dysfunctions, including neurocognitive disturbances, that compromise quality of life during the short life span of patients, one year usually ^1^. These tumors are often resistant to standard treatments which include resection, radiotherapy and chemotherapy with temozolomide ^2^. Numerous studies are focused on new molecular targets to treat GBs ^3-6^; however, none of them has proven effective yet, which is in stark contrast to the considerable progress made in other tumor types. It is therefore necessary to explore new biological concepts that can lead to additional therapeutic strategies against GBs.

The WNT canonical pathway is activated upon the ligand “Wingless-related integration site” (WNT) binding to specific receptors (LRPs and FZD) in the plasma membrane. As a consequence, the destruction complex (APC and Axin) is inactivated and β-catenin (armadillo in *Drosophila*) is released. Further, β-catenin translocates into the cell nucleus where it promotes the expression of target genes (*i.e. Cyclin D1* and *Myc*)^7-8^. The WNT pathway is conserved through metazoans and it plays a central role in brain development ^9^, adult neuronal physiology ^10^ and synaptogenesis ^11^. Perturbations in WNT signaling are associated with neural deficits, Alzheimer’s disease and brain cancer, most notably GB ^12^. WNT and FZD signaling can be deregulated in glioblastoma ^13-14^ (reviewed in ^15^). In particular, one of the hallmarks of bad prognosis is the accumulation of ß-catenin in tumoral cells ^16-17^, indicating an activation of WNT/FZD pathway ^18^.

GB cells extend ultra-long membrane protrusions that interconnect tumor cells ^19^. These tumor microtubes (TMs) are associated with the worst prognosis in molecular subtypes of human gliomas. TMs contribute to invasion and proliferation, resulting in effective brain colonization by GB cells. Moreover, TMs constitute a multicellular network that connects GB cells over long distances, a feature that likely provides resistance against radiotherapy, chemotherapy and surgery ^19-20^. Considering the many cytological similarities of TMs and tunneling nanotubes (TNTs) ^21^, it seems that TMs in aggressive gliomas are the in vivo correlate of TNTs described in cell culture. In addition, TMs seem akin to a basic mechanism of cell-cell connection and molecular communication called “cytonemes” in *Drosophila* ^22^. *Growth Associated Protein-43* (*GAP43*) is essential for the development of TMs and, thus, the tumor cell network which is associated with GB progression ^19^. However, many aspects of this paradigmatic finding in glioma biology are still unexplored, including its impact on neighboring neurons.

Here, we report that *Drosophila* glial cells develop a TM network upon oncogenic transformation, akin to what is known from refined mouse glioma models, and from patients ^19^. These TMs share characteristics in common with *Drosophila* epithelial cytonemes ^22-23^ which are also dependent on igl/*Gap43* expression. TMs relocate Frizzled (Fz1) receptor in the glia-neuron interphase, depleting Wingless (Wg) from neighboring neurons. As a consequence, the number of glioma cells increases, while neuronal synapse number decreases and neurodegeneration ensues. The concept of a Wg/Fz1 signaling equilibrium between glioma cells and neurons is relevant because it redefines GB as a neurodegenerative disease, and because it reveals a potential novel strategy against GB.

## Results

### *Drosophila* glioma network

To study the mechanisms of communication among malignant glial cells and neighboring neurons, we used a previously described *Drosophila* GB model ^24^, which consists in the co-overexpression of constitutively active forms of EGFR (dEGFRλ) and the PI3K catalytic subunit p110α (PI3K92E) driven by the glial specific *repo-Gal4*. This combination stimulates malignant transformation of post-embryonic larval glia, leading to lethal glial neoplasia ^24-25^ which is measured by the increase of glial cell number (GFP^NLS^ positive) compared with a control brain (Figure 1A-C). Based on this *Drosophila* GB model we have evaluated the impact of glial tumor cell proliferation on neighboring neurons.

**Figure 1:**
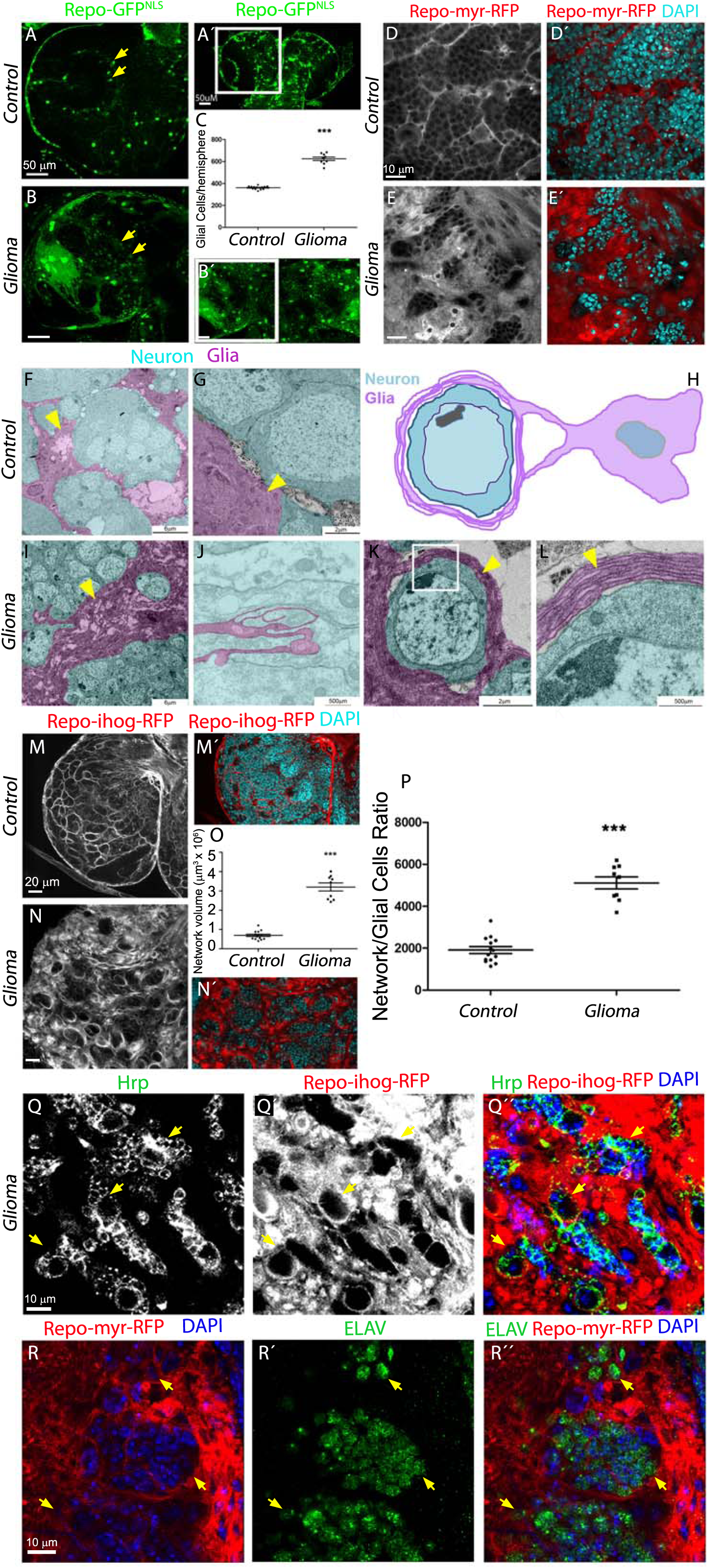
Co-activation of EGFR-Ras and PI3K in *Drosophila* glia causes an expansion of the glial network. Brains from 3rd instar larvae. Glia nuclei are labeled with GFP^NLS^ (green) driven by *repo-Gal4*. Each brain is composed of 2 symmetrical hemispheres. (A-C) In *repo>dEGFR^λ^; dp110^CAAX^*(*glioma*) larvae (B), both brain hemispheres are enlarged and elongated and the number of glial cells is increased relative to *wt* control (A). Lower magnifications are shown in A’ and b’. The quantification of the number of glial cells is shown in (C). Arrows indicate glial nuclei. (D–E) Optical sections of larval brain to visualize the glial network, glial cell bodies and membranes are labeled in red (myrRFP). (D) RFP signal in control brains shows glial somas and the network in *wt* brain. (E) The glioma brain shows a dramatic increase in the membrane projections and in the size of the network. Nuclei are marked with DAPI (blue). (F-L) Transmission electron microscopy (TEM) images of a 3rd instar larval brains expressing HRP in the glial cells. (F-G) HRP deposits label cell profiles membranes in dark, thus identifying glial cells. Coloured images from control brains showing glial cells identified by HRP staining (magenta) and HRP-negative neurons (cyan). (H) Schematic diagram, a glioma cell labeled with HRP (magenta) showing that glioma cells produce a network of TMs that grow to surround neighboring neurons (cyan). (I-L) Several magnifications of Glioma brains showing TMs that grow and enwrap neighboring neurons (cyan). Detail of several layers of a glioma membrane enwrapping a neuron (K-L) and a longitudinal section of a TM (J), arrows indicate glial membranes. (M-P) Glia are labeled with *UAS-ihog-RFP* to visualize active cytoneme/TM structures in glial cells as part of an interconnecting network. (M) In control brains the active glial cytonemes are shown by *repo>ihog-RFP* in red. In glioma brains (N), the TMs grow and expand across the brain, quantification of the network volume (O) and the network/glial cell ratio is shown in P. (Q) Glia is labeled with *UAS-Ihog-RFP* driven by *repo-Gal4* to visualize TMs in glial cells, nuclei are marked with DAPI (blue) and Neurons are stained in green (Hrp) and enwrapped by glial TMs in glioma brains (yellow arrowheads). (R) Glia is labeled with *UAS-myr-RFP* driven by *repo-Gal4* to visualize membrane projections in glial cells, nuclei are marked with DAPI (blue) and Neurons are stained in green (ELAV) and enwrapped by glial TMs in glioma brains (yellow arrowheads). Error bars show S.D. *** P<0.0001. Scale bar size are indicated in this and all figures. Genotypes: (A) w; *repo-Gal4, UAS-GFP^nls^/UAS-lacZ*, (B) *UAS-dEGFR^λ^, UAS-dp110^CAAX^;; repo-Gal4, UAS-GFP^nls^*, (D) w; *Gal80^ts^; repo-Gal4, UAS-myrRFP/UAS-lacZ*, (E, R) *UAS-dEGFR^λ^, UAS-dp110^CAAX^; Gal80^ts^; repo-Gal4, UAS-myrRFP*, (F-G) w; *UAS-HRP:CD2; repo-Gal4/UAS-lacZ*, (I-L) *UAS-dEGFR^λ^, UAS-dp110^CAAX^; UAS-HRP:CD2; repo-Gal4*, (M) w;; *repo-Gal4, UAS-ihog-RFP/UAS-lacZ*, (N, Q) *UAS-dEGFR^λ^, UAS-dp110^CAAX^;; repo-Gal4, UAS-ihog-RFP*

To visualize the total volume of the glial plasma membrane in larval brains, we expressed a myristoylated form of red fluorescent protein (expressed via *UAS-myrRFP*). RFP signal in control brains shows glial soma and the network formed among WT glial cells (Figure 1D, D’). Relative to the control brain, the glioma brain shows a significant enlargement in glial membrane volume (Figure 1E, E’).

In addition, we obtained transmission electron microscopy (TEM) images to visualize glia morphology in control samples (Figure 1F-H). High magnification TEM images show an enlargement of glial cells surface in glioma brain samples (Figure 1I), and the infiltration of glioma cells projections through the brain (Figure 1J). Glioblastoma tumor microtubes (TMs) are described in human GB samples as a cell to cell communication system ^19^, we describe here comparable structures in *Drosophila* glioma brain samples. Magnifications of TEM images show the detail of perineuronal nests of TMs which surround and isolate neurons (Figure 1K-L).

TMs share characteristics with the cytonemes previously described in epithelial *Drosophila* cells ^19^. Ihog (*interference hedgehog*) is a type 1 membrane protein shown to mediate the response to the active Hedgehog (Hh) protein signal ^26^, which accumulates in the epithelial cytonemes ^27^. To visualize glia projections in the entire brain of *Drosophila*, we expressed a *UAS-Ihog-RFP* construct under the control of *repo-Gal4*. This red fluorescent tagged form of Ihog-RFP in epithelial cells labels cellular processes (cytonemes) in the basal region of the wing imaginal discs ^27-28^. The expression of *UAS-Ihog-RFP* under the control of *repo-Gal4* allows the visualization of projections in wild type glial that build an interconnecting network (Figure 1M, M’). The accumulation of Ihog in the cellular projections of transformed glial cells allowed labeling of the TMs-like processes (Figure 1N, N’). We used 3D reconstructions (IMARIS Bitplane, see materials and methods) from confocal stacks of images to quantify the volume of glial membrane projections, and we observed an expansion of TMs in glioma as compared to control brains (Figure 1O). Since the term tumor microtubes (TMs) is the now established terminology to describe thin membrane protrusions from malignant glioma cells in human and murine tumors ^19^, we decided to also use this term from here on for membrane protrusions in GB *Drosophila* cells observed across the brain.

To determine if TMs expand as a consequence of the increase in the number of GB cells, we quantified the volume of the TMs and divided by the number of glial cells (Figure 1P). The results show that TMs volume/glial cell number ratio is higher in GB and therefore we conclude that TMs grow in GB.

A detailed analysis of the contact region between neuron and GB cells revealed that glial TMs wrap clusters of neurons in individual GB “nests” (Figure 1K, L compared with G, and see video S1 and S2), this organization is comparable to previously described perineuronal nests in GB patients ^29^. To confirm the neuronal identity of the cells within the nests, we used antibodies to specifically mark neurons (anti-Hrp and anti-ELAV, green in Figure 1Q-R) and the Ihog-RFP or myr-RFP to visualize the TMs. The results show that TMs infiltrate within neuronal groups and enwrap neurons which segregate neuronal populations.

TMs were previously described to be actin based projections (as cytonemes) ^19^, to further determine the identity of the TMs in *Drosophila* glioma cells, we took advantage of the LifeActin-GFP reporter and observed in confocal images that TMs (ihog positive) are also actin-based (Figure S1A-C’”). Moreover, confocal 3D reconstructions (IMARIS, bitplane) of control and glioma brains with the glial network marked with ihog-RFP and LifeAct-GFP showed that actin based TMs form perineuronal nests (Figure S1A-B’” and Videos S4-S5). Additionally we characterized the TMs with four previously described markers for cytonemes in *Drosophila* ^28^ and orthologues markers of human TMs ^19,28^ (GMA-GFP, GPI-YFP, GFP-MLC and Sqh-GFP in Figure S1D-G). Moreover, we performed functional experiments (Figure S1 H-I”) where the TMs marked with ihog-RFP, are defective after the downregulation of *neuroglian* (*nrg-RNAi*) as it has been previously described in epithelial cytonemes ^30^. All these results together suggests that glioma cells build an organized TMs network around the neurons.

Next we sought to clarify whether similar molecular machineries are involved in human and *Drosophila* TMs. Growth-Associated protein 43 (GAP43) is necessary for TMs formation and function, and drives microtube-dependent tumor cell invasion, proliferation, interconnection, and radioresistance ^19^. We have reproduced this specific and unique characteristic of human GB in the *Drosophila* model. To determine whether the *Drosophila* glial network is susceptible to *GAP43* depletion, as it has been described in human tumor cells ^19^, we knocked down *igloo* (*igl*), the invertebrate GAP-43-like gene ^31^, in glioma cells. Confocal images of larval brains show that the glioma network does not develop upon *igl* silencing and, as a consequence, glial TMs do not enwrap neurons showing a phenotype similar to the control (Figure S1J-L’ and video S1-3).

To exclude the possibility that suppression of TMs expansion and glioma progression was due to a titration of GAL4 activity caused by introducing an additional UAS-transgene, we also tested whether co-expression of *UAS-lacZ* or *UAS-yellow-RNAi* had any rescuing ability in glioma brains. The observed phenotypes such as the number of glial cells and the GB nests were unchanged in the presence of an additional UAS-transgene (Figure S2A-B). This observation indicates that in *Drosophila* glioma igl reproduces the function previously described in human samples.

The genetic disruption of igl/*Gap43* prevents the tumoral network of TMs and halts the overgrowth of glioma cell membranes. Moreover, the direct consequence on flies developing a glioma is larval/pupal lethality. Upon *igl* knockdown, however, the glioma-induced lethality is prevented allowing the emergence of adults (Figure S1M). Interestingly, *igl* knockdown in wild type glial cells neither affects the normal development of neurons and glia, nor their viability (Figure S2C-G). Taking all data together, transformed glial cells take advantage of the igl/Gap43-dependent tumoral network to proliferate and enwrap neurons and, as a consequence, cause death. Thus, the dependency of TMs on igl/GAP43 is conserved between flies and gliomas originated from human tumor cells.

### Wingless signaling in glioma

GB is characterized by the deregulation of many key signaling pathways involving growth, proliferation, survival, and apoptosis, such as p53, pRB, EGFR, PI3K, STAT3 or WNT ^32-33^. WNT signaling has long been suggested as a hallmark in gliomagenesis associated with the proliferation of stem-like cells in human GBs ^34^. WNT signaling promotes tumoral cell proliferation and dissemination as well as resistance to chemo and radiotherapy (reviewed in ^15,35^). To assess the prevalence of WNT genes/pathway deregulation in GB, we searched for mutations related to WNT pathway in the Collection of Somatic Mutations in Cancer (COSMIC). We analyzed data from 922 samples of grade IV astrocytomas (GB) searching for gain of expression or duplications. First, we analyzed *Wnt* family genes, which encode the ligands of the WNT pathway. In particular, we analyzed WNT6 that is related to the self-renewal ability of GB cells ^36^, WNT3A and WNT1 overexpression that is detected in glial stem cells ^37^ and WNT5A and WNT2 that were shown to induce migration of GB cells ^38-39^. Our analysis of the COSMIC database revealed that WNT1, WNT3A, WNT6 or WNT5A genes do not appear mutated in GB samples and only 6 cases showed mutations in WNT2 (0,6%) (Figure S3A). WNT signaling is a hallmark of GBs, but as no evident changes in WNT expression were found, we searched for mutations in FZD genes, that encode the receptors of WNT. In particular, we analyzed FZD2 and FZD9 that are linked to self-renewal ability in GB ^36,39-40^ and FZD4, a positive WNT regulator, which is a causative effector for invasive phenotypes of GB cells ^41^. We did not find any case of a GB patient with a gain of expression in FZD2 or FZD4, and barely 4 cases (0,4%) showed a mutation for FZD9. The total number of mutations related with WNT or FZD genes accounts for 5% of the total GB samples analyzed (Figure S3A). In addition, we analyzed data from GB in The Cancer Genome Atlas (TGAC) for WNT ligands, FZD receptors and transcriptional targets of WNT pathway databases (Figure S3B-D) through the Xena Functional Genomics Explorer (https://xenabrowser.net/). We analyzed expression levels in primary GB tissue and non tumoral tissue, transcriptional targets for WNT pathway (Figure S3B) are upregulated in GB samples. However, WNT ligands from the canonical WNT pathway are not upregulated (Figure S3C) and among FZD receptors, only FZD7 shows significant changes in GB tissue (Figure S3D). Taking all these data together, in spite of WNT pathway activation in GB tissue, there is no correlation between overexpression of WNT ligands and GB development.

Nevertheless, previous reports indicate that WNT targets, such as FoxM1 which promotes nuclear localization of β-catenin, or β-catenin itself are related with Glial Stem Cells (GSC) maintenance and tumorigenesis. Indeed, they are standard biomarkers for GB bad prognosis ^16,42-45^. β-catenin activation (translocation to the nuclei) is a downstream event of WNT pathway. It has been identified in 19% of surgical samples from adult GB patients and in 30% of surgical samples from pediatric patients. Moreover, WNT inhibition leads to suppression of tumor growth, proliferation in cultures, and a modest induction of cell death ^34^.

Given the discrepancy between the presence of WNT pathway markers and the lack of mutations in GB samples, we decided to study *Wingless* (*wg*) expression, the fly homologue to human WNT, in our glioma model. We quantified anti-Wg signal intensity in glial cells and neuronal tissue and obtained the Wg in glia vs Wg in neurons pixel intensity ratio. The results showed that Wg protein is homogeneously distributed in control larval brains with a slight increase (1,5 ratio) in glial membranes (Figure 2A-A’’, C) but glioma brains showed a four-fold increase in accumulation of Wg in tumoral cells (Figure 2B-B”, C), in line with WNT accumulation in human GBs ^38,46^. To determine whether Wg could be signaling to the glial cells, we assessed the presence of Frizzled (Fz) receptors in glial membranes. Monoclonal antibodies were used to visualize the Fz1 and Fz2 receptors in control samples, the quantification of anti-Fz1 signal in glia vs neurons shows that Fz1 receptors are localized homogeneously across the brain with a small accumulation in glial membranes (Fz1, Figure 2D-D”, H). Fz2 staining shows the expected pattern in the wing disc ^47^ as a control (Figure S4A) and it is localized homogeneously in control brains S4B-B’”). However in glioma samples, Fz1 is highly accumulated in glial membranes (Figure 2E-E”, H). No changes for Fz2 were detected in glioma brains (Figure S4C-C’”). In addition, a detailed analysis of glioma brains revealed that Fz1 protein specifically accumulated in TMs (iHog+) (Figure 2E”). The results indicate that Fz1 is preferentially accumulated in perineuronal nests of TMs, in contact with neighboring neurons. These data suggest that the glioma TMs network contributes to Wg/Fz1 signaling.

**Figure 2:**
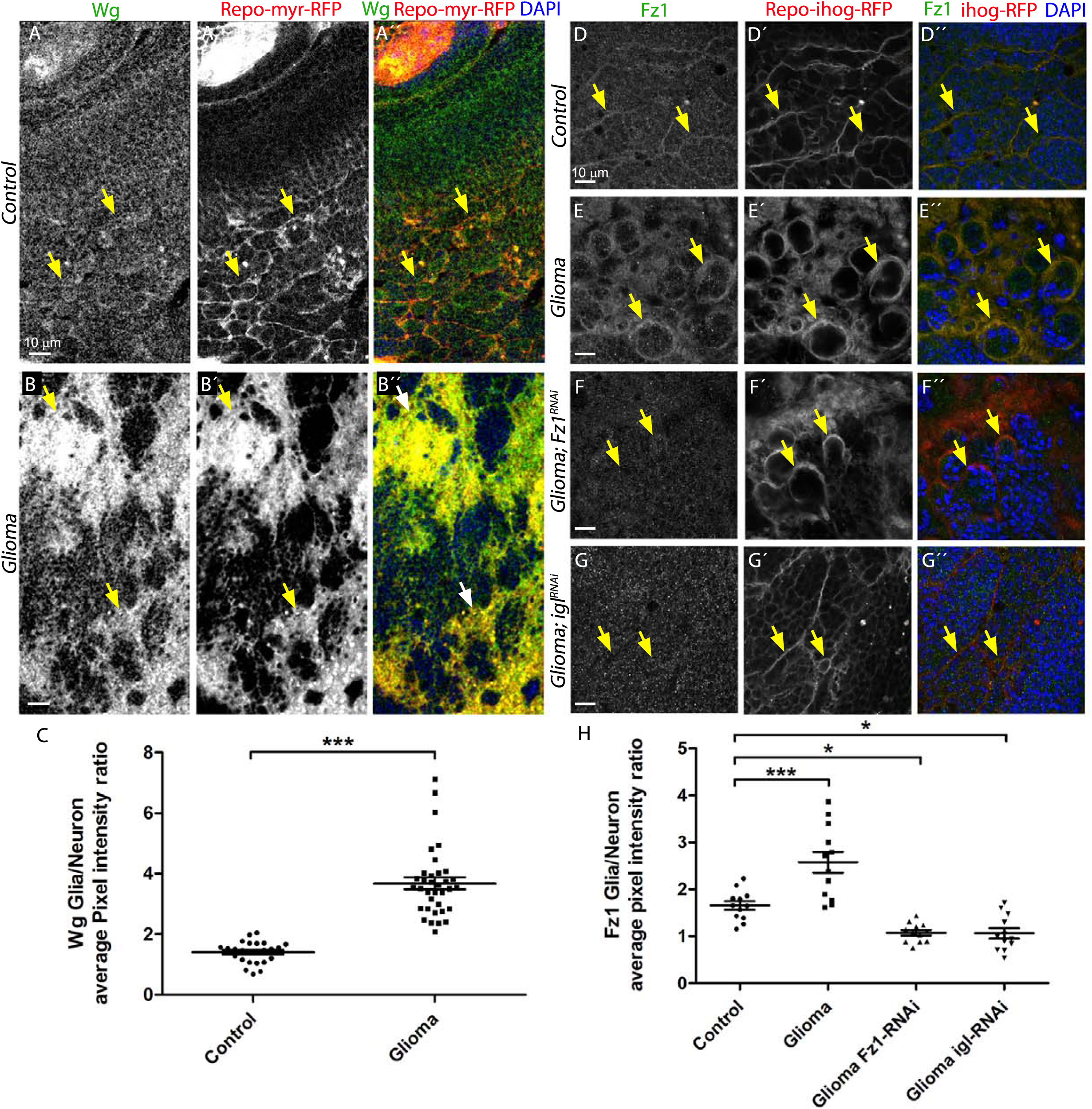
Wingless/Fz1 accumulate in glioma cells. Larval brain sections with glial cell bodies and membranes labeled in red (myrRFP) and stained with Wg antibody show homogeneous expression in the control brains (A) in green. In the glioma brains Wg accumulates in the glial transformed cells (B), the Wg average pixel intensity ratio between Glia/Neuron quantification is shown in (C). Arrows indicate Wg staining in glial membranes. (D-F) Glial cells are labeled with *UAS-Ihog-RFP* to visualize the glial network, and stained with Fz1 (green). (D) Fz1 is homogeneously distributed in control brains, with a slight accumulation in the Ihog+ structures. (E) Fz1 accumulates in the TMs and specifically in the projections that are in contact with the neuronal clusters. (F) Upon knockdown of *fz1* in glioma brains, the tumoral glial network is still formed but Fz1 is not detectable. (G) Knockdown of *igl* in glioma brains restores a normal glial network and Fz1 shows a homogeneous distribution along the brain section. Arrows indicate Fz1 staining in glial membranes. (H) Fz1 average pixel intensity ratio between Glia/Neuron quantification ^−^. Nuclei are marked with DAPI. Error bars show S.D. * P<0.01 *** P<0.0001. Genotypes: (A) w; *Gal80^ts^; repo-Gal4, UAS-myrRFP/UAS-lacZ*, (B) *UAS-dEGFR^λ^, UAS-dp110^CAAX^; Gal80^ts^; repo-Gal4, UAS-myrRFP*, (C) w;; *repo-Gal4, ihog-RFP/UAS-lacZ*, (D) *UAS-dEGFR^λ^, UAS-dp110^CAAX^;; repo-Gal4, UAS-ihog-RFP*, (E) *UAS-dEGFR^λ^, UAS-dp110^CAAX^; UAS-Fz1-RNAi; repo-Gal4, UAS-ihog-RFP*, (F) *UAS-dEGFR^λ^, UAS-dp110^CAAX^;; repo-Gal4, UAS-ihog-RFP/UAS-igl-RNAi*

To evaluate the contribution of Fz1 to the progression of the glioma, we knocked down the Fz1 receptor (using *UAS-Fz1-RNAi*) in transformed glial cells. First we validated *UAS-Fz1-RNAi* tool in epithelial tissue from larvae wing imaginal discs. Even though Fz1 expression in wing imaginal disc is discrete, upon *UAS-Fz1-RNAi* expression in the posterior compartment (marked with GFP), there is a reduction in Fz1 protein signal compared with the anterior compartment from the same tissue (Figure S5A). Brains showed a significant reduction of Fz1 protein expression (Figure 2F, H). However, TMs network was still formed (Figure 2F’). Furthermore, we inhibited the development of the glioma network by expressing *UAS-igl-RNAi* and stained the brains for Fz1. Under these conditions the network does not expand, Fz1 does not accumulate in TMs and it shows a homogeneous distribution comparable between glia and neurons (Figure 2G-G’, H). These results indicate that Fz1 accumulation in TMs is a consequence of the glioma network but suggests that Fz1 accumulation in the TMs is dispensable for glioma network formation.

### Glial Fz1 interacts with neuronal Wg

The abnormal distribution of Wg and Fz1 in glioma brains could be due to either an increase in gene expression or to redistribution of the protein. First, to determine *wg* and *Fz1* transcription levels in glioma, we performed quantitative PCR experiments from brain RNA extracts. The results showed that *wg* and *Fz1* transcription levels are comparable in control and gliomas (supp. Figure S6A). To consider the mRNA translation or protein stability and degradation, we measured total Fz1 and Fz2 protein levels in Western blot experiments. Control and glioma brain protein extracts were blotted and incubated with anti-Fz1 or Fz2 antibodies; Tubulin was used as a loading control (Supp. Figure S6B). Quantification of the membranes showed no significant changes for Fz1 protein levels in glioma (Supp. Figure S6B). Moreover, we performed *in situ* hybridization experiments to detect Wg and Fz1 mRNA expression in control and glioma brains. The results are consistent with the qPCRs results (Figure S6A) showing no differences for *wg* or *Fz1* transcription between controls and gliomas (Figure S6C).

The data indicate that in spite of a higher signal for Wg and Fz1 proteins in glioma cells by immunofluorescence, there are no changes in gene expression. We hypothesized that Fz1 is transported and accumulated along glioma TMs, which contact neighboring neurons and receive Wg from them.

According to this hypothesis, the glial membranes would be in close proximity to neuronal membranes to allow Fz1-Wg interaction. To assess this, we performed GRASP experiments ^48^. This technique determines intimate physical interaction among glia and neurons, in the range of 20-40nm (synaptic distance) which is compatible with protein-protein interaction ^48^. Each split fragment of the green fluorescent protein (GFP) was expressed in neurons (*elav-lexA*) or glial (*repo-Gal4*) cells, respectively (see Material and Methods). Only upon intimate contact between the two split fragments GFP fluorescence is reconstituted. Control samples (Figure 3A-C”) showed a discrete signal corresponding to the physiological interaction between glia and neurons, nevertheless, upon glioma induction a massive GFP signal from GRASP reporter was detected (Figure 3D-F”). This result indicates that, in a glioma condition, there is a significant increase of glia-neuron membrane interaction, consistent with the TEM images (Figure 1H-L) showing perineuronal nests of TMs which surround and isolate neurons.

**Figure 3:**
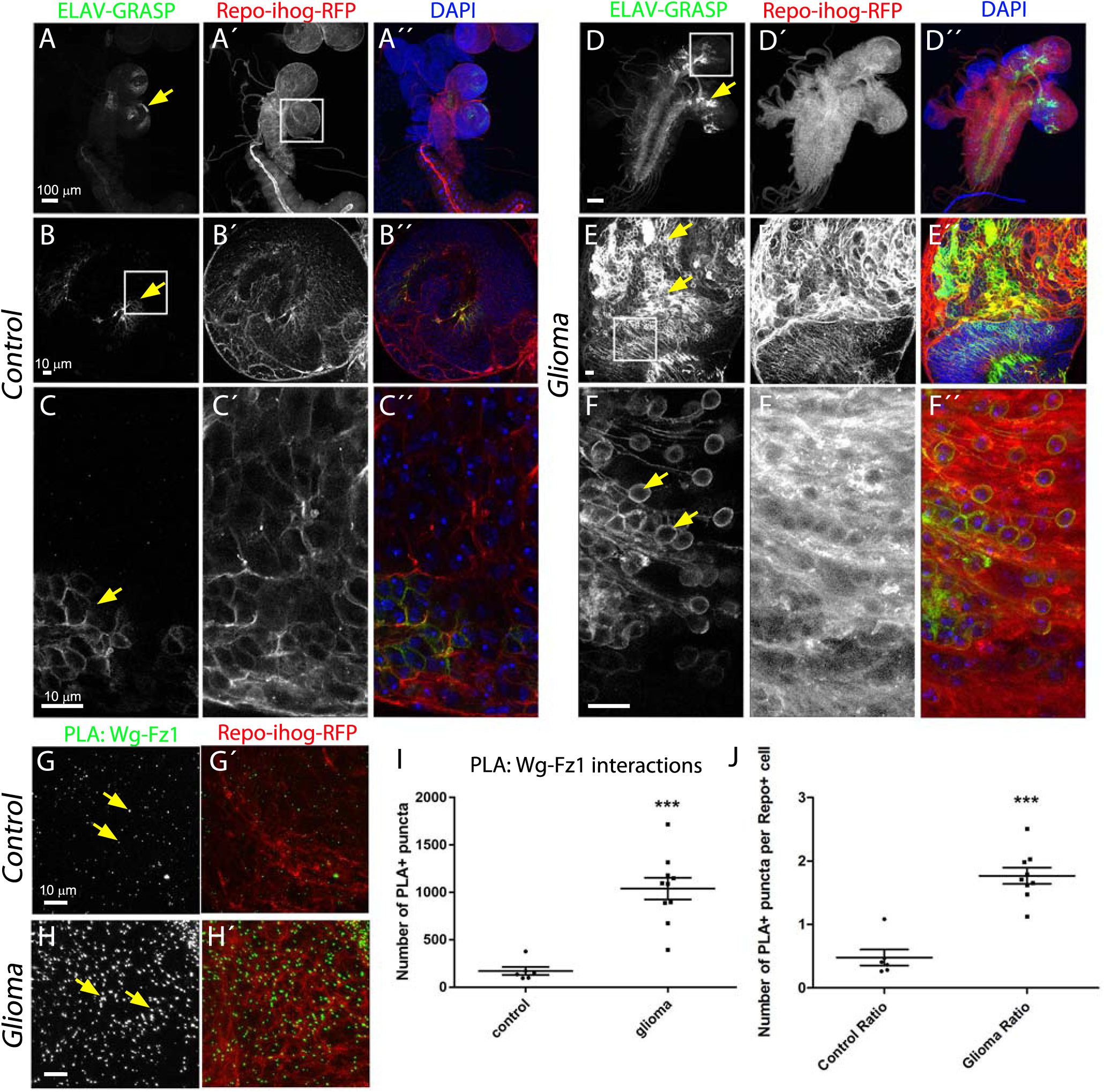
Fz1 in glia interacts with neuronal Wg. GRASP technique was used and both halves of green fluorescent protein tagged with a CD4 signal to direct it to the membranes (CD4-spGFP) were expressed in neurons (*elav-lexA*) and glial (*repo-Gal4*) cells respectively. Only upon intimate contact GFP protein is reconstituted and green fluorescent signal is visible. (A-C) Control brains showed a discrete signal corresponding to the physiological interaction between glia and neurons. (D-F) In glioma brains a massive GFP signal from GRASP reporter is detected. Arrows indicate GRASP reconstitution GFP signal. (G-H) Proximity ligation assays (PLA) were performed in control and glioma brains to quantify the interactions between Wg and Fz1. (G) Control brains showed a discrete number of puncta (green) showing the physiological interactions. (H) Glioma brains showed a five-fold increase in the number of puncta, quantified in (I). The number of PLA+ Wg-Fz1 interactions per Repo+ cell in control and glioma brains is shown in (J). Nuclei are marked with DAPI (blue). Arrows indicate PLA^+^ puncta. Error bars show S.D. *** P<0.0001. Genotypes: (A-C) w;; *elav-lexA; repo-Gal4, UAS-ihog-RFP/UAS-CD4-spGFP1-10, lexAop-CD4-spGFP11*, (D-F) *UAS-dEGFR^λ^, UAS-dp110^CAAX^; elav-lexA; repo-Gal4, UAS-ihog-RFP/UAS-CD4-spGFP1-10, lexAop-CD4-spGFP11*, (G) w;; *repo-Gal4, ihog-RFP/UAS-lacZ*, (H) *UAS-dEGFR^λ^, UAS-dp110^CAAX^;; repo-Gal4, UAS-ihog-RFP*

We had observed specific signal localization for Wg and Fz1 in glioma membranes (myr-RFP/ihog-RFP) (Figure 2B-B”, E-E”). To confirm the physical interaction of Fz1 and Wg proteins we performed proximity ligation assays (PLA) (see Materials and Methods) ^49^. This quantitative method reports the interactions between two proteins with a resolution of 40nm ^50^. Control brains showed a discrete number of puncta showing the physiological interactions (Figure 3G-H’ and quantifications in 3I, J)). However, glioma brains showed a five-fold increase in the number of PLA^+^ puncta (Figure 3H’,I, J) indicating a significant increase in the number of Fz1-Wg interactions. Moreover, we calculated the number of Wg-Fz1 interactions per glial cell in control and glioma brains and the results show that the number if interactions per cells is higher in the glioma brains (Figure 3J). These results confirm that glioma cells accumulate Fz1 receptor in TMs, and then this specific receptor interacts with Wg. But Wg is not upregulated in the glioma brain so we hypothesize that Wg comes from neighboring neuronal membranes and it relocates and accumulates in glioma cell membranes.

### Wg/Fz1 pathway targets are active in glioma and inactive in neurons

Wg targets are indicators of Wg/Fz1 activity in the recipient cell. Armadillo/βCatenin is a cytoplasmic protein which, upon activation of Wg pathway, translocates into the nucleus and activates transcription of target genes ^12,51^. To determine if Fz1 is signaling in gliomas as a consequence of Wg-Fz1 interaction, we stained control and glioma brains for Armadillo (Arm) with a specific antibody that detects Arm protein and is mainly visible in its cytoplasmic inactive form (Cyt-Arm) ^52^. Cyt-Arm was homogeneously distributed along the brain in control samples (Figure 4A, D). However, glioma brains showed accumulation of Cyt-Arm in the cytoplasm of the neurons and a reduction in glioma cells (Figure 4B, D). This result suggests that, in glioma brains, Wg/Fz1 pathway is inactive in neurons and active in glioma cells. More importantly, this data shows that the network expansion and the accumulation of Fz1 in the TM projections have an effect on neighboring neurons. Further, we dismantled the glioma network expressing *UAS-igl-RNAi* or downregulated Fz1 (*UAS-Fz1-RNAi*) and stained for Cyt-Arm. Under this condition, Cyt-Arm was homogeneously distributed along the brain similar to the control (Figure 4C, D and S7A) demonstrating that the network is required to promote Wg signaling in the transformed glia.

**Figure 4:**
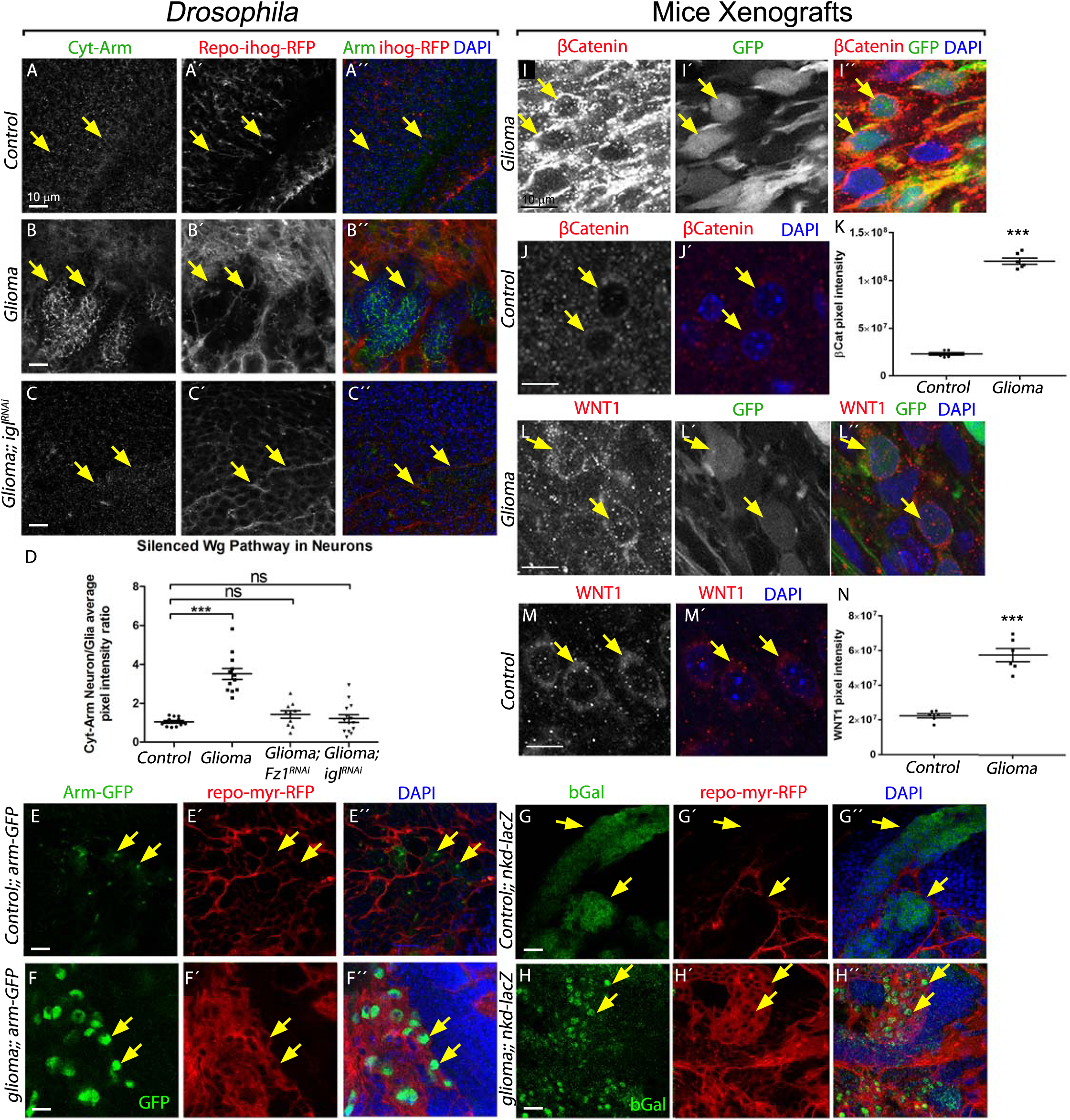
Wg signaling pathway is active in glioma cells, and the glioma inactivates it in neuronal clusters in glioma brains. Larval brain sections with glial cytonemes labeled in red and stained with Arm (green). (A) Cytoplasmic-Armadillo (Cyt-Arm) is homogeneously distributed in control sections. (B) In glioma brains Cyt-Arm accumulates in the neurons cytoplasm where it is inactive. (C) Knockdown of *igl* in glioma brains restores a normal glial network and Arm does not accumulate showing a homogeneous distribution similar to the control. Arrows indicate Cyt-Arm staining at the Glia-neuron interphase. (D) Cyt-Arm average pixel intensity ratio between Neuron/Glia quantification showing the Wg signaling pathway silencing in neurons in a glioma brain. Glial cell bodies and membranes are labeled with myrRFP (red). (E-H) Wg signaling pathway reporters *arm-GFP* (E-F) and *nkd-lacZ* (stained with anti-bGal (G-H) in control and glioma brains show activation of the pathway in glioma cells compared with the reporter activation mostly in neurons in the control brains. Arrows indicate cells with reporter activation. (I-N) Confocal immunofluorescence single plane images of S24 GBSC NMRI nude mice brains (glioma) and NMRI nude mice (control) brains stained with human anti-βCatenin (I-J) and WNT1 (L-M) both show in grey (red in the merged image) an increase in the glioma samples. The corresponding quantification of the pixel intensity is shown in K and N. Green signal from tumor cell GFP expression allows specific detection of S24 GBSC related structures in the mouse brain (I’, L’). Arrows indicate glioma or control cells. Nuclei are marked with DAPI. Error bars show S.D. *** P<0.0001 and ns for non-significant. Genotypes: (A) w;; *repo-Gal4, ihog-RFP/UAS-lacZ*, (B) *UAS-dEGFR^λ^, UAS-dp110^CAAX^;; repo-Gal4, UAS-ihog-RFP*, (C) *UAS-dEGFR^λ^, UAS-dp110^CAAX^;; repo-Gal4, UAS-ihog-RFP /UAS-igl-RNAi*, (E) *w; Gal80^ts^; repo-Gal4, UAS-myrRFP/arm-GFP*, (F) *UAS-dEGFR^λ^, UAS-dp110^CAAX^; Gal80^ts^; repo-Gal4, UAS-myrRFP/ arm-GFP*, (G) *w; Gal80^ts^; repo-Gal4, UAS-myrRFP/nkd04869a-lacZ*, (H) *UAS-dEGFR^λ^, UAS-dp110^CAAX^; Gal80^s^; repo-Gal4, UAS-myrRFP/nkd04869a-lacZ*

To further confirm that Wg pathway is active in the glial transformed cells and silenced in neurons, we used five additional Wg pathway reporters, namely *arm-GFP, nkd-lacZ, tsh-lacZ, fz4-GFP* and *dally-LacZ* ^54-55^. The results showed that all these reporters were active in glioma cells and inactive in neurons compared to control brains (Figure4E-H and Figure S7B-G”) which confirm that the Wg/Fz1 pathway is inactive in neurons confronted to a GB, and active in GB cells.

Since we observed specific signal localization and accumulation of Wg and Fz1 in glioma membranes and consequently the activation of the pathway in glioma cells, we wondered if these results were conserved in human glioma cells. Therefore, we used a primary patient-derived glioblastoma culture xenograft model using S24 cells kept under stem-like conditions (see Materials and Methods), which reproduce previously described Scherer modes, perivascular migration and spread ^29,56^. GFP-marked S24 cells were injected in the brain of NMRI nude mice, and brains were removed and analyzed 90 days later (as previously described in ^19^). To validate our data, we performed a series of more diffuse tumor parts (history grade II-like tumor periphery where normal brain has just been colonized) and denser tumor parts (grade III-IV like, from central areas) GB images from S24 xenografts brain sections.

We stained the samples for β-catenin and WNT1 proteins in S24 xenograft brain sections and compared them to control samples (Figure 4I-N and Figure S8G-H). We observed a significant increase of both proteins in GB cells, in line with previous observations. These data indicate that WNT1 is deposited in GB cells and activates WNT pathway; consequently β-Catenin is upregulated to mediate GB cells malignancy ^16^. Moreover, images of the WNT1 and βCatenin staining of grade II and III GB images from S24 xenografts brain sections and quantification of the pixel intensity indicates that the accumulation of WNT1 and activation of βCatenin correlates with the growth of the tumor (Figure S8A-F).

To determine Fz1’s contribution to the proliferation of *Drosophila* gliomas we quantified the number of transformed glial cells upon *Fz1* downregulation. A specific Repo antibody was used to mark glial cell nuclei in the brains. The data show a significant increase of glial cell number in glioma brains, which is prevented by *Fz1* downregulation or TM network dismantlement (Figure 5A-E). In addition, we studied the adult survival by quantifying the number of flies with glioma and/or *Fz1-RNAi* expression. First, Fz1 downregulation in normal glial cells allows 80% adult survival. On the other hand, the glioma caused 100% lethality in larval or pupal stages. By contrast, glioma lethality was reverted upon *Fz1* downregulation in repo cells, and 70% of the animals reached adulthood (Figure 5F). In conclusion, Fz1 is not necessary for the glioma network overgrowth but Wg/Fz1 signaling is necessary for the increase in the number of tumoral cells and the associated lethality.

**Figure 5:**
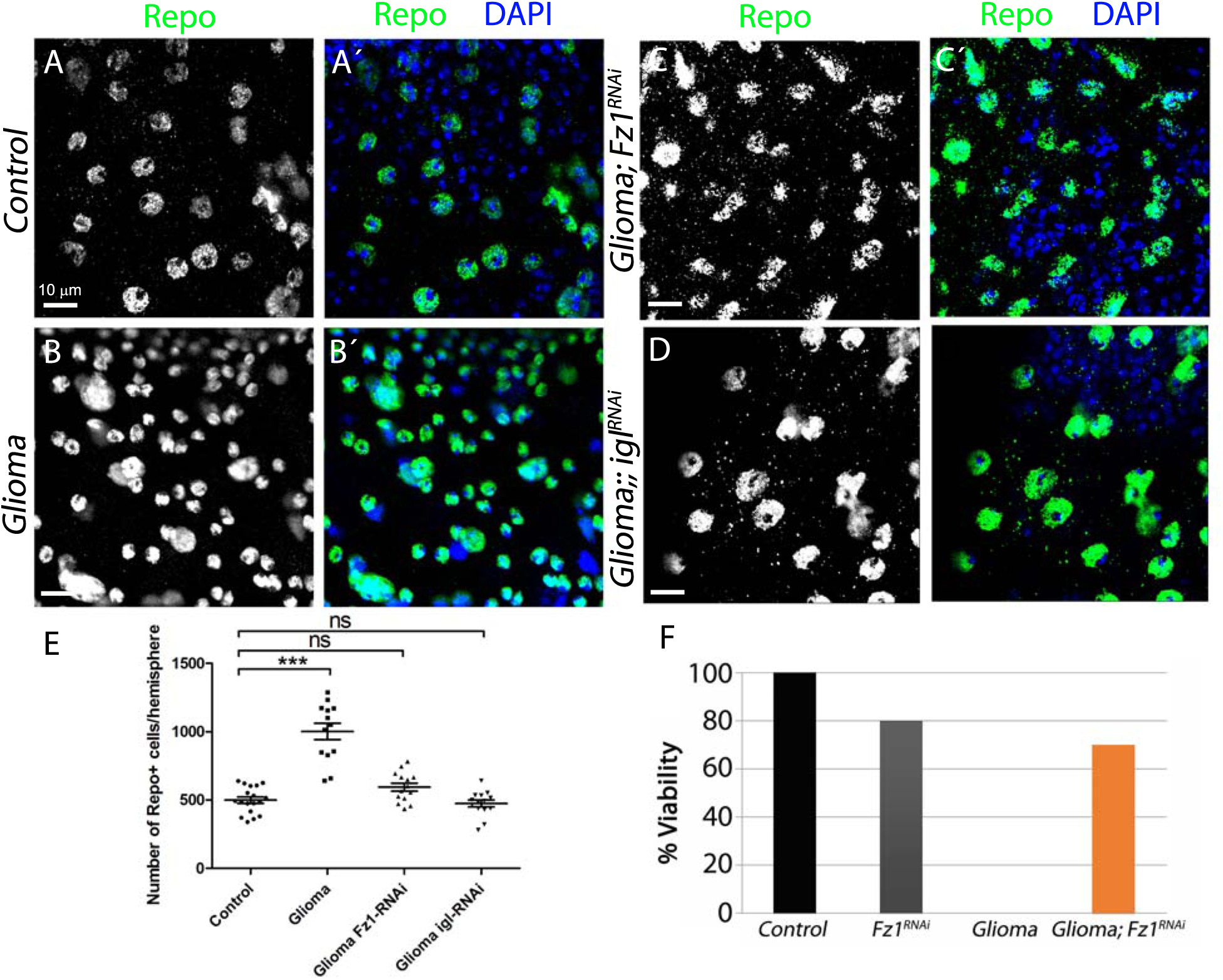
Glioma network is responsible for the increase in the number of glial cells. Larval brain sections with glial cell nuclei stained with Repo (green). The number of glial cells is quantified in the following genotypes: (A) Control, (B) Glioma showing an increase in Repo+ cells. (C) Upon knockdown of *Fz1* in glioma brains, the number of glial cells is partially restored (D) knockdown of *igl* in glioma brains restores the number of glial cells similar to the control. (E) Quantification of the number of Repo+ cells. Nuclei are marked by DAPI (blue). (F) Viability assay showing the % lethality induced by the glioma that is partially rescued upon knockdown of *fz1*. Error bars show S.D. *** P<0.0001, and ns for non-significant. Genotypes: (A) w;; *repo-Gal4, ihog-RFP/UAS-lacZ*, (B) *UAS-dEGFR^λ^, UAS-dp110^CAAX^;; repo-Gal4, UAS-ihog-RFP*, (C) *UAS-dEGFR^λ^, UAS-dp110^CAAX^; UAS-Fz1-RNAi; repo-Gal4, UAS-ihog-RFP*, (D) *UAS-dEGFR^λ^, UAS-dp110^CAAX^;; repo-Gal4, UAS-ihog-RFP /UAS-igl-RNAi*, (F) 1. w;; *repo-Gal4, ihog-RFP/UAS-lacZ 2*. w; *UAS-fz1-RNAi; repo-Gal4, UAS-ihog-RFP 3.UAS-dEGFR^λ^, UAS-dp110^CAAX^;; repo-Gal4, UAS-ihog-RFP 4. UAS-dEGFR^λ^, UAS-dp110^CAAX^; UAS-fz1-RNAi; repo-Gal4, UAS-ihog-RFP*

### Wg/Fz1 pathway disruption in adult brains

To discard that these results are restricted to a developmental process, we performed similar experiments in adult brains. We included the *tub>Gal80^TS^* system to repress Gal4 activation during all developmental stages (see Materials and Methods), glioma was induced 4 days after adult eclosion when adult brain is fully differentiated. Analysis of adult flies survival indicate that animals with glioma start dying at day 9 after the glioma induction until the 14th day, significantly earlier that control flies. (Figure S9A). 7 days after glioma induction we stained for Cyt-Arm, Fz1 and Wg control and glioma adult brain samples and quantified (n>15) the average pixel intensity ratio (Glial cells membranes RFP+ vs neuronal tissue (Figure S9B-H). The results reproduce the findings from larval brains: Fz1 and Wg proteins are accumulated in glioma tissue and Cyt-Arm signal shows an increase of wg-pathway activity in glioma cells. Therefore, there are no differences between larval and adult brains regarding Wg-pathway in glioma cells and neurons. Moreover, we analyzed *Drosophila* adult negative geotaxis behavior (climbing assay), as an indication for possible motor defects associated with neurodegeneration. The results showed symptoms of neurodegeneration in glioma flies compared to controls (Video S6).

### Gliomas cause Neurodegeneration

Previous results suggest that a neurodegeneration process is taking place in glioma brains. To determine whether the glioma is causing neurodegeneration we quantified the number of active zones (synapses) in the neuromuscular junction (NMJ). This well-established system has been used for decades to study neurodegeneration in *Drosophila* ^57-60^, motor neuron soma is located in the central nervous system, but synapse counting can be accurately done in synaptic buttons located in adult or larva muscular wall. Adult NMJs Quantification of confocal images stained with anti-bruchpilot (Nc82) revealed a significant reduction in the number of synapses in glioma adult (Figure S9I) or larval NMJs compared to controls (Figure 6A-B, F) and therefore, a neurodegenerative process, which is prevented by Fz1 downregulation or TM network dismantlement (igl downregulation) (Figure 6C-D, F).

**Figure 6:**
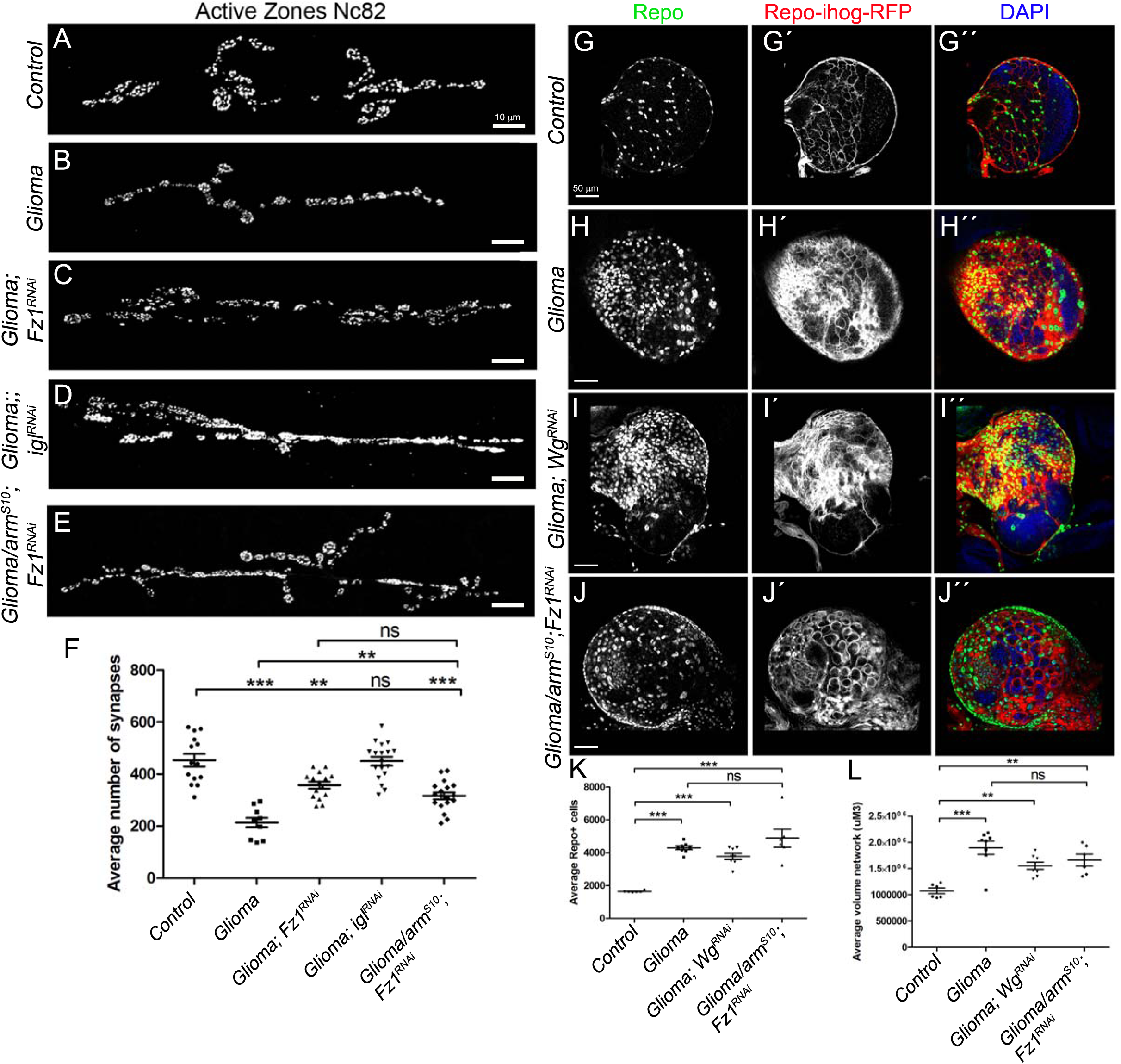
Gliomas cause Neurodegeneration and Wg expression in glioma cells is dispensable for tumor progression. Neurons from the larval neuromuscular junction are stained with Nc82 showing the synaptic active zones. (A-F) Upon glioma induction (B) the number of synapses (grey) is reduced when compared with the control (A). The number of synapses is restored upon knockdown of *Fz1* (C), *igl* (D) or *armS10; Fz1-RNAi* (E). The quantification of synapse number in all genotypes is shown in (F). (G-L) *wg* knockdown (I) in glioma cells (*wg-RNAi*) or *armS10; Fz1-RNAi* (J) does not prevent glioma cell numbers increase nor glioma TMs volume expansion quantified in (K-L). Error bars show S.D. *** P<0.0001, ** P<0.001 and ns for non-significant. Genotypes: (A, G) w;; *repo-Gal4, ihog-RFP/UAS-lacZ*, (B, H) *UAS-dEGFR^λ^, UAS-dp110^CAAX^;; repo-Gal4, UAS-ihog-RFP*, (C) *UAS-dEGFR^λ^, UAS-dp110^CAAX^; UAS-Fz1-RNAi; repo-Gal4, UAS-ihog-RFP*, (D) *UAS-dEGFR^λ^, UAS-dp110^CAAX^;; repo-Gal4, UAS-ihog-RFP /UAS-igl-RNAi*, (E, J) *UAS-arm^S10^/UAS-dEGFR^λ^, UAS-dp110^CAAX^; UAS-Fz1-RNAi; repo-Gal4, UAS-ihog-RFP*, (I) *UAS-dEGFR^λ^, UAS-dp110^CAAX^; UAS-wg-RNAi; repo-Gal4, UAS-ihog-RFP*

### Wg expression in glioma cells is dispensable for tumor progression

Fz1 receptor is accumulated in glioma TMs and contributes to Wg depletion and pathway equilibrium disruption. To determine whether the source of Wg is neuronal or glial, we silenced *wg* expression in neurons exposed to glioma, or in glioma cells. First we validated *UAS-wg-RNAi* tool in epithelial tissue, wing imaginal discs. Activation of *UAS-wg-RNAi* in the posterior compartment (marked with GFP) cause a reduction of specific monoclonal anti-wg (Figure S5B). Pan-neuronal *wg* silencing (*elav>wg RNAi*) is lethal in line with our hypothesis regarding the requirement of Wg in neuronal biology. Besides, *wg* knockdown (*wg-RNAi*) in glioma cells (*Repo>PI3K; EGFR; wg-RNAi*) does not prevent glioma cell number increase (Figure 6G-I, K) nor glioma TMs volume expansion (Figure 6G-I, L), these results suggest that *wg* expression in glioma cells is not relevant for glioma progression.

Moreover, to stress on the contribution of Fz1 receptor as mediator for Wg depletion, we generated glioma cells and silenced *Fz1* expression, in addition we expressed a constitutively active form of *armadillo* (*UAS-armS10*) ^61^ to activate Wg pathway downstream Wg-Fz1 in these gliomas (Figure 6E-F and J-L). We counted the number of synapses and observed that it is comparable to glioma + *Fz1RNAi* (Figure 6C-F). This result suggests that the reduction in the number of synapses is specifically mediated by Fz1 accumulation. To confirm that the Fz1 depletion and Arm signaling produces a glioma like condition we counted the number of glial cells and TM network volume (Figure 6G-L). We observed that in this case the number of glioma cells increased and TMs expanded (Figure 6J-L) similar to a glioma (Figure 6H).

### Wg/Fz1 pathway disruption causes neurodegeneration

Neuronal development and physiology are dependent on Wg/Fz1 signaling and disruptions in this signaling pathway lead to synapse loss, an early symptom of neurodegeneration (reviewed in ^62-65^. To determine if an imbalance in Wg distribution caused by glioma cells can affect the neighboring neurons, we aimed to determine the contribution of Fz1/Wg pathway to neuronal physiology. To inhibit Wg/Fz1 pathway signaling we expressed *UAS-fz1-RNAi* or *UAS-wg-RNAi* in motor neurons under the control of a *D42-Gal4* driver ^66^ and quantified the number of active zones (synapses) in the neuromuscular junction (NMJ). Quantification of confocal images stained with anti-bruchpilot (Nc82) revealed a significant reduction in the number of synapses and therefore, a neurodegenerative process (Figure 7A-D). These data suggest that Wg/Fz1 signaling pathway activity in neurons is necessary in for synaptogenesis.

**Figure 7:**
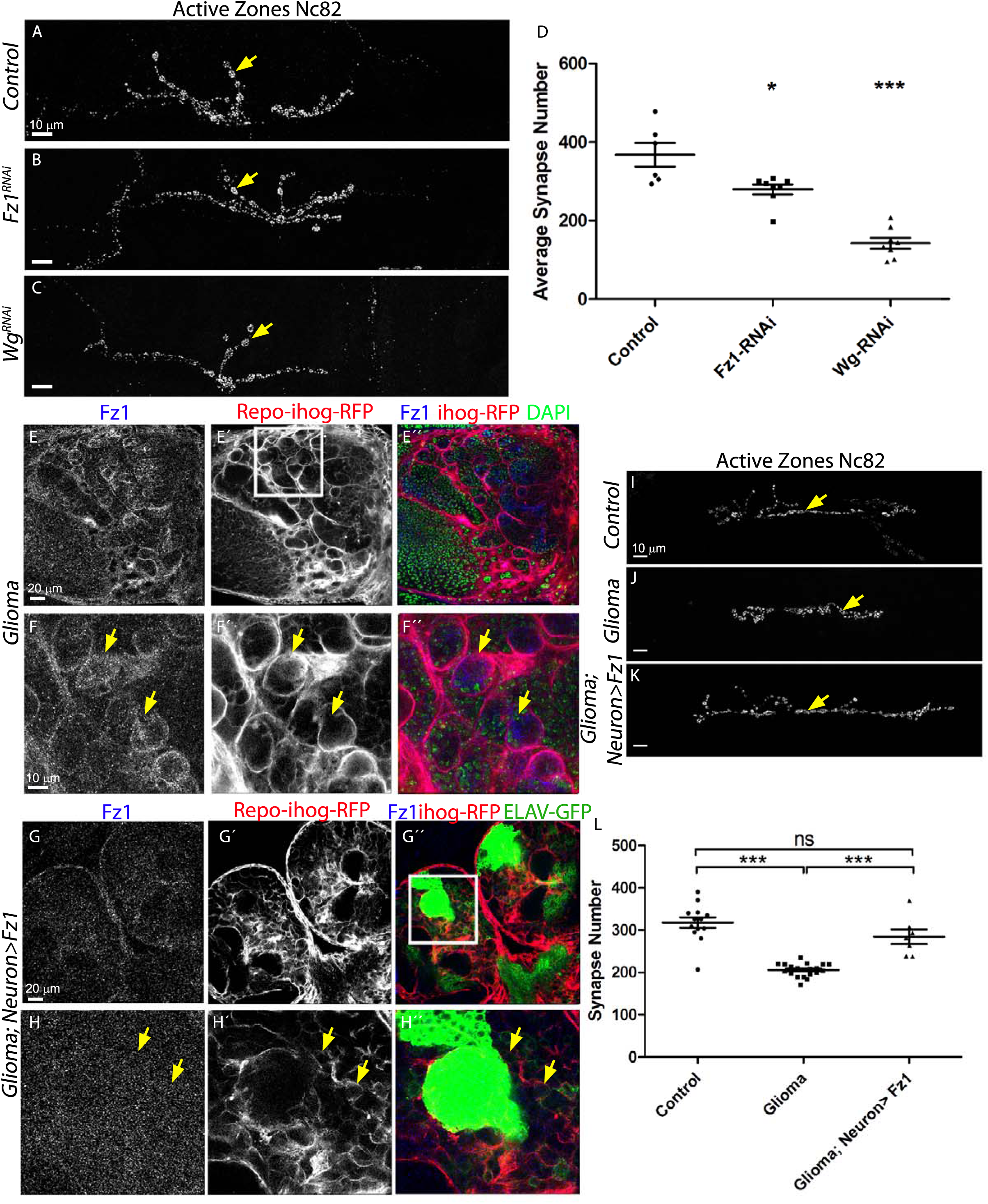
Knockdown of the Wg signaling pathway results in neurodegeneration and restoration of the glia-neuron Wg/Fz1 signaling equilibrium inhibits glioma progression. Neurons from the larval neuromuscular junction are stained with Nc82 showing the synaptic active Zones. (A-D) Upon knockdown of *wg* (C) or *Fz1* in D42 neurons (B) the number of synapses (grey) is reduced when compared with the control (A). Arrows indicate synapses. The quantification of synapse number in all genotypes is shown in (D). (E-H) Larval brain sections with glial network labeled with *UAS-Ihog-RFP* in red and stained with Fz1 (grey or blue in the merge). Neurons are labeled with *lexAop-CD8-GFP* driven by *elav-lexA*. Fz1 overexpression in neurons (green) restore homogeneous Fz1 protein distribution (blue) in the brain, rescue brain size and neuron distribution (G and magnification in H) compared to (E and magnification in F) where the *elav-lexA* is not present in the glioma brains, Nuclei are marked with DAPI in (E-F) (green). Arrows indicate Fz1 staining in the glial membranes at the Glia-neuron interphase of glioma brains and its restored localization in G-H. (I-L) Neurons from the larval neuromuscular junction are stained with Nc82 showing the active Zones. Upon glioma induction the number of synapses (grey) is reduced (J) when compared with the control (I). The number of synapses is restored upon overexpression of Fz1 specifically in the neurons (K). Arrows indicate synapses. The quantification of synapse number is shown in (L). Error bars show S.D. *** P<0.0001, * P<0.01 or ns for non-significant. Genotypes: (A) w; *UAS-CD8-GFP; D42-Gal4/UAS-lacZ*, (B) w; *UAS-CD8-GFP/Fz1-RNAi; D42-Gal4*, (C) w; *UAS-CD8-GFP/wg-RNAi; D42-Gal4*, (E-F, J) *UAS-dEGFR^λ^, UAS-dp110^CAAX^; lexAop-Fz1; repo-Gal4, UAS-ihog-RFP*, (G-H, K) *UAS-dEGFR^λ^, UAS-dp110^CAAX^; lexAop-fz1/elav-lexA, lexAop-CD8-GFP; repo-Gal4, UAS-ihog-RFP*, (I) w; *repo-Gal4, ihog-RFP/UAS-lacZ*

So far, we have demonstrated that glioma cells cause a disruption of Wg/Fz1 signaling in neurons (Figure 4) and our data suggest that this is dependent on Fz1 accumulation in glioma cells (Figure 2). Next, we decided to restore this signaling equilibrium by overexpressing Fz1 receptor in neurons surrounded by glioma cells. To avoid crossed expression we generated Fz1 transgenic flies based on the LexA-LexAop expression system ^67^, which is independent of the Gal4-UAS system used to generate the glioma. We validated this newly generated tool in *Drosophila* brains. *LexAop-Fz1* was activated in neurons under the control of *elav-LexA* and monoclonal anti-Fz1 showed higher signal in neurons and functional activity revealed by anti-Arm staining (Figure S5C-E). Oversized glioma brains showed the expected glioma network compartmentalizing neurons in the brain (Figure 7E-F”). However, Fz1 overexpression in neurons restored homogeneous Fz1 protein distribution in the brain (Figure 7G, H), rescued brain size (Figure 7G-G”) and neuron distribution and morphology (Figure 7H’-H”). In addition, Fz1 equilibrium restoration partially rescued lethality and most animals reached adulthood. To verify Fz1 activation of the pathway we stained for Wg and Cyt-Arm (Figure S10). As previously shown, glioma brains showed a heterogeneous distribution for Wg protein (Figure S10A-A”) and, as a consequence, an imbalance in pathway activation reported by Cyt-Arm accumulation (Figure S10 C-C”). As expected, neuronal Fz1 overexpression in glioma brains restored Wg distribution and Cyt-Arm signal to a control situation (Figure S10B-B”, D-D”)

To further determine the effect of Wg/Fz1 signal restoration in neurons, we quantified the number synapses in their NMJs. Neuronal morphology is disrupted by glioma (Figure 7I, J) and restored by Fz1 overexpression in neurons neighboring glioma cells (Figure 7J, K). Moreover, synapse number reduction upon glioma induction is restored by Fz1 overexpression in neurons (Figure 7L). All these results indicate that the Wg/Fz1 pathway disruption caused by glioma is responsible of the synapse loss. Restoration of the signaling equilibrium between glia and neurons prevents synapse loss and therefore, neurodegeneration.

### Glioma depletes Wg from neuronal membranes

The actual mechanisms of Wingless delivery have been under debate. This protein was initially described as a diffusible secreted protein ^68^. Recent studies have proved that Wg secretion is not necessary for *Drosophila* development ^69^. A membrane-tethered version of Wg protein ^70^ (*Wg^NRT^*) can substitute the endogenous gene, mimic Wg normal functions and produce viable organisms ^69^. We took advantage of this tool to determine the cellular mechanisms mediating glioma Wg retrieval from neurons. We created a genetic combination to substitute one copy of endogenous Wg with one copy of *wg^NRT^* exclusively in neurons (by using the LexAop system). In addition, upon LexA system activation, neurons are marked with membranous GFP (CD8-GFP) while the rest of the cells are wild type. Moreover, this Wg^NRT^ is tagged with an HA which allows to discriminate from the endogenous Wg protein. We induced a glioma and marked the glial cells in red (ihog-RFP) (see materials and methods and Figure 8G). First, we studied normal control brains that express Wg^NRT^ exclusively in neurons in the absence of glioma. We stained with anti-HA and observed a positive signal both in neurons and glial cells but not in the control without the elav-LexA driver, indicating that the HA signal is not a false positive (Figure 8A-B, H). Finally, we performed the experiment in the glioma model to examine the interaction of glioma cells with Wg^NRT^ expressing neurons (Figure 8E-F-I) and stained with anti-HA. The images show a homogeneous signal for HA throughout the brain in glia and in neurons (See figure H-I magnifications for more detail), suggesting that glioma cells can deplete this non-secretable/membrane tethered version of Wg from neurons. We have quantified the number of HA^+^ puncta in glia and neuron from control and glioma samples. The results suggest that glial cells are able to sequester membrane anchored Wg^NRT^ from neuronal membranes at comparable rates under both physiological and glioma conditions (Figure 8J). However, there was a detectable reduction in the number of glioma cells, similar to previous results when Wg/Fz1 equilibrium in glial vs neurons was restored (Figure 7C, E and Figure 7G-H”). Since Wg^NRT^ is anchored to neuronal membranes, it would be expected to reduce the total Wg signaling in glioma cells thereby decreasing cell proliferation/survival, thus resulting in a rather normal sized brain (Figure 8A, C, E). Moreover, heterozygous *wg^NRT^/wg* prevented glioma network progression (Figure 8D’vs F’). In conclusion, glioma cells produce a network TMs that reach neighboring neurons, increasing intimate membrane contact that facilitates neuronal-Wg sequestering mediated by the Fz1 receptor in glioma TMs. Since Wg/Fz1 signaling in glioma mediates cell number and tumor progression, targeting this cellular interaction may be a new candidate for future therapies.

**Figure 8:**
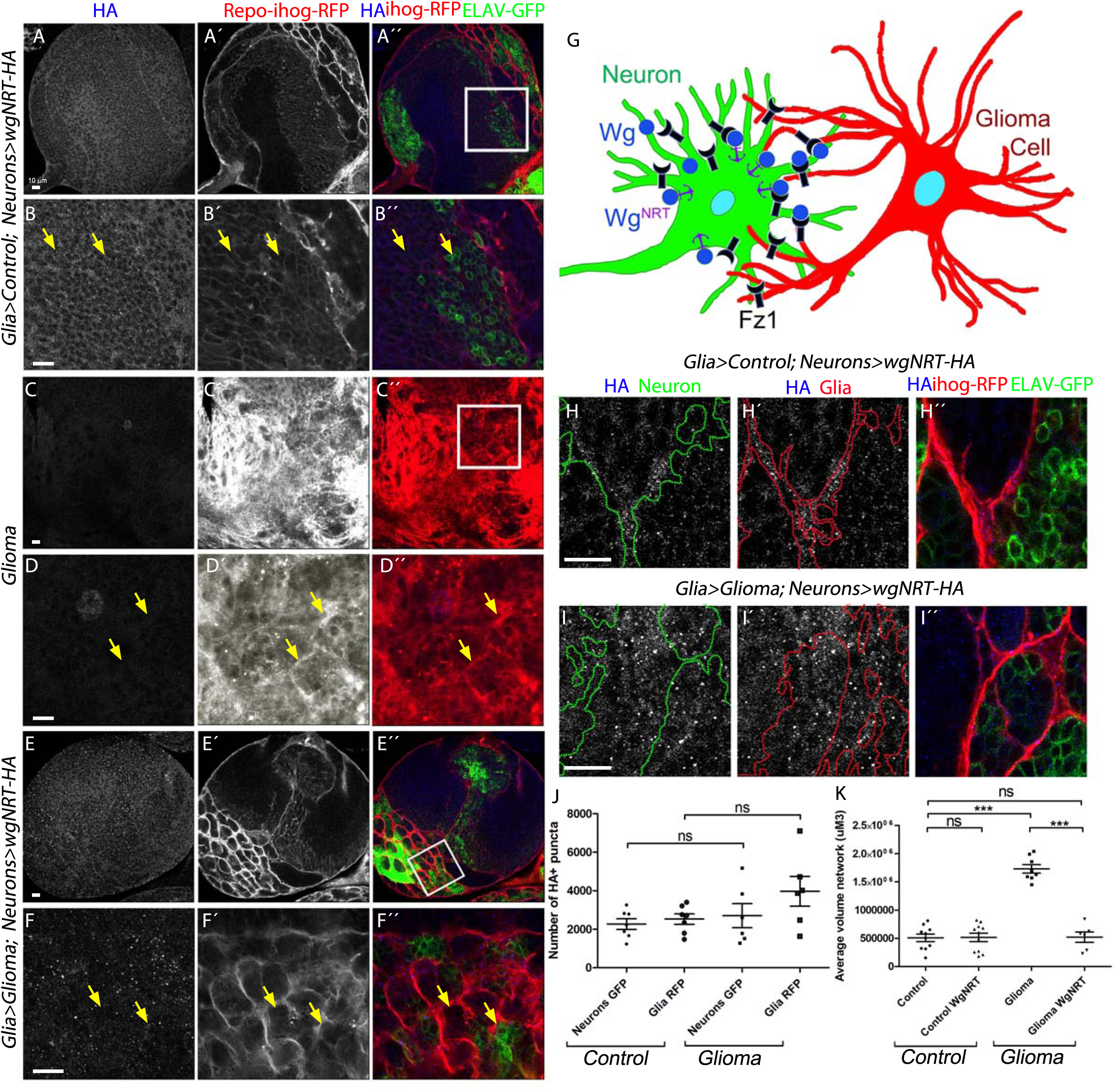
Glioma depletes Wg from neuronal membrane. Larval brain sections with glial network labeled with *UAS-Ihog-RFP* in red and stained with HA (blue). (A, magnification in B, H) Control brains express Wg^NRT^-HA and the anti-HA (grey or blue in the merge) staining shows a positive signal in both neurons (green) and glial cells (red). (C-D) Glioma samples that do not express Wg^NRT^ in neurons do not show HA nor GFP signal. (E, magnification in F, I) Glioma brains with membrane anchored Wg (Wg^NRT^-HA grey or blue in the merge) in neurons (green), show a homogeneous signal for HA (grey or blue in the merge) in both glioma cells and neurons, quantified in (J), and the glial network volume size is restored in these animals (K). Arrows indicate HA^+^ staining in glial or neuronal membranes. Error bars show S.D. *** P<0.0001, ns for non-significant. (G) Schematic diagram of this experiment: a neuron labeled with GFP (green) and a glioma cell labeled with ihog-RFP (red) showing that glioma cells produce a network of TMs that grow to reach neighboring neurons. Intimate membrane contact facilitates neuronal-Wg (blue) sequestering mediated by Fz1 receptor (black) from glioma. In this experiment neurons express a membrane anchored version of Wg (Wg^NRT^ represented as Wg in blue with a purple anchor) which is more difficult for the glioma to retrieve from the neuron. Genotypes: (A-B, H) w; *>wg>wgNRT-HA, PaxRFP/ elav-lexA, lexAop-CD8-GFP; repo-Gal4, UAS-ihog-RFP/lexAop-flp*, (C-D) *UAS-dEGFR^λ^, UAS-dp110^CAAX^; >wg>wgNRT-HA, PaxRFP; repo-Gal4, UAS-ihog-RFP/lexAop-flp*, (E-F, I) *UAS-dEGFR^λ^, UAS-dp110^CAAX^; >wg>wgNRT-HA, PaxRFP/ elav-lexA, lexAop-CD8-GFP; repo-Gal4, UAS-ihog-RFP/lexAop-flp*

## Discussion

In addition to cell autonomous features of tumor cells, including founder mutations, recent evidences indicate that microenvironment signals contribute to glioma progression. Neuroligin-3 (NLGN3) is a synaptic protein cleaved and secreted after neuronal activity which promotes PI3K-mTOR signaling stimulating glioma growth. Thus, NLGN3 mediates an autocrine/paracrine loop in glioma cells which perpetuates tumoral features ^71^ (reviewed in ^72^). Also, neural precursor cells (NPC) from the subventricular zone (SVZ) produce chemoattractants (SPARC/SPARCL1, HSP90B and pleiotrophin) which facilitate glioma invasion of the SVZ through Rho/ROCK signaling ^73^.

We showed recently that TM network formation determines GB tumor malignancy, confers radiotherapy resistance and influences patient’s prognosis ^19^. TMs stability in GB is sensitive to *GAP43* expression in tumoral cells ^19^. Also, *Tweety homologue-1* (*TTHY1*) expression in GB cells, mediated by NLGN3 regulates TM formation ^74^.

This study shows that TMs intercalate among neurons and enwrap them in perineuronal nests ^29^ structures establishing an intimate link glioma-neuron. Then GB cells make direct contact via TMs and deprive neurons of WNT.

Expression data from human cancer databases indicate that glioma cells do not upregulate *Wnt* expression, neither upregulate its receptors, however downstream targets of the pathway such as β-catenin are upregulated. Instead, the results in the fly model show that glioma cells relocate Fz1 receptor in the TMs allowing depleting Wg from neurons. Thus we use the term vampirization as the action of GB cells to exhaust or prey upon healthy cells (neurons) in the manner of a vampire, as they drain Wg/WNT and cause the demise of the neurons. Consistent with these data, in the patient-derived GB xenograft model, where WNT1 is deposited in GB cells and the WNT pathway is activated, β-catenin is upregulated. The available data suggest that GB TMs grow towards the source of Wg. However, as TMs expand upon Fz1/Wg signaling, the question regarding the exact order of events remains open. Do TMs require some initial stimuli from the source of Wg to grow? Alternatively, do TMs initiate growth triggered by glial internal signals and directed through a gradient of neuron-secreted attractants?

Concerning the mechanism of Wg vampirization, we have expressed a non secretable, HA-tagged version of membrane-tethered Wg^NRT^ in neurons. In that experiment Wg is detected within glial cells demonstrating that GB cells can take Wg directly from the neuronal membrane. However, further studies are still required to determine the precise mechanism of neuronal Wg depletion by GB cells TMs.

It is widely observed that brain tumors and related ailments can cause cognitive decline and neuronal dysfunction (reviewed in ^75^). High-grade glioma patients continue to display cognitive deficits after surgery, radiotherapy or chemotherapy ^76-78^. The most common deficits concern memory, executive functions and general attention beyond the effects of age, education and gender ^79^. Nevertheless, the molecules mediating neuronal degeneration need to be determined.

Synapse loss is an early step in neurodegeneration ^80-82^ which is consistent with the cognitive defects observed in GB patients. Nonetheless, cognitive defects can be observed also in patients with excess of synapses as in the case of fragile X syndrome ^83-84^. GB cells can stimulate aberrant synapses associated with seizures ^85^ which are compatible with cognitive dysfunctions. Neuronal death is a later event in neurodegenerative processes such as Alzheimers’s disease ^86-88^. In GB patients, neuronal death is under debate and this issue should be addressed with further data. There is preclinical work ^88-89^ supporting Glioma-induced neuronal death due to glutamate cytotoxicity, in addition clinical studies from ^90^ support neuronal death in GB patients. However, it is certainly very difficult to draw clear conclusions from clinical samples or clinical courses, considering that therapy, antiepileptics and the pure space occupation plus the edema contribute to the neuronal dysfunction, degeneration and cell death.

In particular, neuronal cell loss is typically found at and around glioblastomas, and neurocognitive disturbances are a frequent finding in glioma patients. Although evidences from our experience and from neuropathology expertise, this is an open debate which requires further attention.

The data also reveal that reestablishing Wg signaling equilibrium by Fz1 overexpression in neurons, not only restores synapse number but also blocks GB progression. Functional disruption of the equilibrium between GB glia and neurons is described here for the first time. Possibly, this mechanism could be valid for other molecules related to tumor progression such as Notch, Hedgehog or TGF. Moreover, cytoneme-like structures play a role in development and in other cell types ^91-93^. Hence, we propose that cytoneme-like structures in physiological conditions and TMs in pathological GB conditions could redistribute limited amounts of signaling molecules among competing cell types, therefore long range redistribution of signaling molecules could be a general mechanism for the cells to compete for different resources.

This study integrates for the first time the oncogenic nature of glioma with the neuronal degeneration caused by Wg depletion. This innovative concept of glioma-induced neurodegeneration opens the possibility of combined treatments to fight GB progression and associated neurodegeneration at the same time. Our data demonstrate that making the neurons more competitive for secretable factors such as Wg already has an impact in GB tumor growth, although it remains to be demonstrated what type of drugs can carry out these actions.

The rapid transformation of GB cells and the heterogeneity of mutations in these tumors are a handicap for genetic therapies and monoclonal therapies. In our view, cellular features such as the network shared by GB cells emerge as an alternative to tackle tumor progression. Among the possible new strategies, TMs dynamic and cellular transport of receptors to the TMs could be a target to prevent GB proliferation and neurodegeneration. GAP43 has emerged as a functional component of GB network formation. Recent studies indicate that other proteins such as Flotilin ^28,94^, participate in cytoneme dynamics. The discovery of molecules regulating TM/cytoneme biology arises as potential targets for cancer treatment.

## Supporting information

## Author contributions

Conceptualization, S.C.T. M.P. and FW; Methodology, S.C.T., VR, M.P., M.L.P., E.S. and N.F.; Investigation, S.C.T., FW, VR, M.P. M.L.P, E.S and N.F.; Writing – Original Draft, M.P, FW and SCT.; Writing – Review & Editing, S.C.T. and M.P.; Funding Acquisition, S.C.T and FW.; Supervision, S.C.T.

## Acknowledgements

We thank Professor Alberto Ferrús, Professor Helena Richardson, Dr. Paco Martín, Dr. Elena Santana, Patricia Jarabo and anonymous reviewers for critiques of the manuscript and for helpful discussions. Clemencia Cuadrado for fly stocks maintenance. We want to thank JF de Célis and C. Martínez Ostalé for their critical help with *in situ* hybridizations. We are grateful to R. Read, I. Guerrero, M. Milan, A. Baena-López, E. Martín-Blanco, David Strutt, the Vienna *Drosophila* Resource Centre, the Bloomington *Drosophila* stock Centre and the Developmental Studies Hydridoma Bank for supplying fly stocks and antibodies, and FlyBase for its wealth of information. We acknowledge the support of the Confocal Microscopy unit and Molecular Biology unit at the Cajal Institute and the *Drosophila* Transgenesis Unit at CBMSO for their help with this project. MP holds a fellowship from the Juan de la Cierva program IJCI-2014-19272 and SCT holds a contract from the Ramón y Cajal program RYC-2012-11410 from the Spanish MICINN. Research has been funded by grant BFU2015-65685P. Authors declare no conflicts of interest.

## Experimental Procedures

### Fly stocks

Flies were raised in standard fly food at 25°C.

Fly stocks from the Bloomington stock Centre: *UAS-GFP^nls^* (BL4776), *UAS-lacZ* (BL8529), *UAS-myr-RFP* (BL7119), *UAS-igl-RNAi* (BL29598), *arm-GFP* (BL8555), *nkd04869a-lacZ* (BL25111), *D42-Gal4* (BL8816), *GFP-fz1-GFP* (BL59780), *repo-Gal4* (BL7415), *puc-lacZ, UAS-CD8-GFP* (BL32186), *tub-gal80^ts^* (BL7019), *elav-lexA (BL52676), lexAop-CD8-GFP* (BL32205), *lexAop-flp* (BL-55819), *UAS-armS10 (BL4782), sqh-GFP (BL57145), UAS-CD4-spGFP1-10, lexAop-CD4-spGFP11* (obtained from BL58755), *en-Gal4, UAS-GFP (from BL25752)*. Fly stocks from the Vienna *Drosophila* Resource Centre: *UAS-fz1-RNAi* (v105493), *fz4-GFP* (v318152), *UAS-wg-RNAi* (v104579), *UAS-yellow-RNAi* (v106068), *UAS-nrg-RNAi* (v107991). *GFP-sls (MLC, ZCL2144 from http://flytrap.med.yale.edu), UAS-dEGFR^λ^, UAS-PI3K92E* (dp110^CAAX^) (A gift from R. Read), *UAS-ihog-RFP* (a gift from I. Guerrero), *tsh-lacZ* and *dally-lacZ* (gifts from M. Milan), *lexAop*-fz1 (generated in this study), *FRT Wg FRT NRT–Wg-HA, pax –Cherry (a gift from A. Baena-López), lifeactin-GFP* (a gift from I. Guerrero), for electron microscopy studies, we used the *UAS-HRP:CD2* as reporter ^95^, *UAS-GPI-YFP ^96^, UAS-GMA-GFP ^97^*.

### *Drosophila* glioblastoma model

The most frequent genetic lesions in human gliomas include mutation or amplification of the Epidermal Growth Factor Receptor (EGFR) gene. Glioma-associated EGFR mutant forms show constitutive kinase activity that chronically stimulates Ras signaling to drive cell proliferation and migration ^98-99^. Other common genetic lesions include loss of the lipid phosphatase PTEN, which antagonizes the phosphatidylinositol-3 kinase (PI3K) signaling pathway, and mutations that activate PI3KCA, which encodes the p110a catalytic subunit of PI3K ^98-99^. Gliomas often show constitutively active Akt, a major PI3K effector. However, EGFR-Ras or PI3K mutations alone are not sufficient to transform glial cells. Instead, multiple mutations that coactivate EGFR-Ras and PI3K/Akt pathways are required to induce a glioma ^29^. In Drosophila, a combination of EGFR and PI3K mutations effectively causes a glioma-like condition that shows features of human gliomas including glia expansion, brain invasion, neuron dysfunction, synapse loss and neurodegeneration ^24,100-101^. Moreover, this model has proved to be useful in finding new kinase activities relevant to glioma progression.^25^ To generate a glioma in *Drosophila* melanogaster adult flies, we used the Gal4/UAS system ^102^ as described above (*repo-Gal4>UAS-EGFRλ,UAS-dp110*. To restrict the expression of this genetic combination to the adulthood, we used the thermo sensitive repression system Gal80^TS^. Individuals maintained at 17°C did not activate the expression of the UAS constructs, but when flies were switched to 29°C, the protein Gal80^TS^ changed conformation and was not longer able to bind to Gal4 to prevent its interaction with UAS sequences, and the expression system was activated.

### Generation of Transgenic flies

*LexAop-Frizzled1* construct was generated by RECOMBINA S.L. Fz1 *(CG17697)* CDS was synthesized by overlapping g-block assembly. The complete 17665bp fragment was amplified using the high fidelity Phusion taq polymerase (Thermo fisher Scientific) and *Eco.Friz.Fw* and *Xba.Friz.Rv* primers. PCR amplicon was cloned in pJET entry vector (Thermo Fisher Scientific), then Frizzled fragment was released with *EcoRI/XbaI* restriction enzymes and sub-cloned into destination *pLOTattB* plasmid.

*Eco.Fz1.Fw* 5’-GAATTGGGAATTCATGTGGCGTCAAATCCTG-3’ *Xba.Fz1.Rv* 5’-TCTAGACTAGACGTACGCCTGCGCCC-3’

Transgenic flies were injected and *Frizzled1* fragment was inserted in the chromosome 2L by the *Drosophila* microinjection service (CBMSO-CSIC) using the following stock: *y[1] M{vas-int.Dm}ZH-2A w[*]; M{3xP3-RFP.attP’}ZH-22A* (BL24481). Transgenic flies were selected individually by eye color (w+) and balanced with *CyO*.

### Antibodies for Immunofluorescence

Third-instar larval brains, were dissected in phosphate-buffered saline (PBS), fixed in 4% formaldehyde for 30min, washed in PBS + 0.1 or 0.3% Triton X-100 (PBT), and blocked in PBT + 5% BSA.

Antibodies used were: mouse anti-Wg (DSHB 1:50), mouse anti-Repo (DSHB 1:50), mouse anti-Fz1 (DSHB 1:50), mouse anti-Fz2 (DSHB 1:50), Rabbit anti-Fz1 ^103^, 1:300), mouse anti-Arm (DSHB 1:50), mouse anti-β-galactosidase (Sigma, 1:500), rabbit anti-GFP (Invitrogen A11122, 1:500), mouse anti-GFP (Invitrogen A11120, 1:500), mouse anti-Nc82 (DSHB 1:20), mouse anti-ELAV (DSHB 1:50), Rabbit anti-Hrp (Jackson Immunoresearch 111-035-144, 1:400), mouse anti-HA (12CA5 Roche 11583816001 1:100), rat anti-HA (Roche 11867423001, 1:200).

Secondary antibodies: anti-mouse Alexa 488, 647, anti-rabbit Alexa 488, 647 (Thermofisher, 1:500). DNA was stained with 2-(4-amidinophenyl)-1H-indole-6-carboxamidine (DAPI, 1μM).

### Cell culture, fixation and histology of S24 Xenograft model

The S24 cell line was derived as a primary glioblastoma culture (Lemke et al., 2012; Osswald et al., 2015). For the S24 glioma model, 50.000 S24:GFP cells (stably transduced by lentivirus) were injected into the cortex in 8-10 week old male NMRI nude mice (Charles River, Sulzfeld, Germany, n=2). Cells were cultivated under serum-free conditions in DMEM-F12 as sphere cultures (Thermo Fisher Scientific Inc., Waltham, MA, USA) supplemented with 2% B-27 (Thermo Fisher Scientific Inc., Waltham, MA, USA), 5 μg/ml human insulin (Sigma-Aldrich Corporation, St. Louis, MO, USA), 12.8 ng/ml heparin (Sigma-Aldrich), 0.4 ng/ml EGF (R&D Systems Inc., Minneapolis, MN, USA) and 0.4 ng/ml FGF (Thermo Fisher Scientific Inc., Waltham, MA, USA). Animals were sacrificed 90 days after glioma cell injection with age-matched wild-type NMRI nude mice (n=2) which were used as control.

Brains were fixed with transcardial perfusion with 40 ml PBS and 40 ml 4 % PFA. The brain was removed and postfixed in 4 % PFA for 4 hrs at room temperature. Afterwards the brains were stored in PBS at 4 °C in the dark.

For histology, S24:GFP tumor-bearing brains were coronally cut on a vibratome (Sigmann Elektronik, Hüffenhardt, Germany) into 100 μm sections. The sections were permeabilized with 1 % TX100 for 3 hrs and counterstained with primary antibodies against beta-catenin (abcam, ab32572) and WNT1 (abcam, ab15251) for 3 hrs in 0.2% TX100 and 5 % FCS. Sections were washed three times with 0.2% TX100 and 5 % FCS and counterstained with secondary antibodies couples to Alexa-647 and Alexa-546 (Invitrogen) as well as DAPI for 3 hrs. The sections were washed three times in 1 × PBS, pH=7.4, and mounted on coverslips using self-made moviol. Images were acquired on a confocal laser-scanning microscope (Leica SP8, Leica, Germany) using a × 63 immersion oil objective (numeric aperture=1.4). z-Stacks were acquired with a pixel size of 141 nm and 300-nm z-steps.

All animal experiments were approved by the regional animal welfare committee (permit number: G132/16 Regierungspräsidium Karlsruhe).

### Western blots

For western blots, we used NuPAGE Bis-Tris Gels 4–12% (Invitrogen), and the following primary antibodies: mouse anti-Fz1 (DSHB 1:500), mouse anti-Fz2 (DSHB 1:500) and mouse anti-tubulin (1:10,000 Sigma), we use Tubulin as a loading control instead of actin because the tumor microtubes are Actin positive and tubulin negative as previously described ^19^. There were 3 biological replicates and Relative Fz1 Average pixel intensity was measured using measurement tool from Image Studio Lite Ver 5.2 and normalized against Tubulin.

### Proximity ligation assay

DUO92101 Duolink^®^ In Situ Red Starter Kit Mouse/Rabbit with DUO92013 Duolink In Situ Detection Reagents FarRed (Sigma).

The interaction between Wg and Fz1 in Drosophila larval brains was detected *in situ* accordingly to the instructions of the manufacturer. Briefly, primary antibody incubation against Wg (mouse anti-Wg (DSHB 1:50) and Fz1 (Rabbit anti-Fz1 ^103^, 1:300)) were applied using the same conditions as immunocytochemistry staining. Duolink secondary antibodies against the primary antibodies were then added. These secondary antibodies were provided as conjugates to oligonucleotides that were able to form a closed circle via base pairing and ligation using Duolink ligation solution when the antibodies were in close proximity ^49^ at a distance estimated to be <40 nm. The detection of the signals was conducted by rolling circle amplification using DNA polymerase incorporating fluorescently labeled nucleotides into the amplification products. The resulting positive signals were visualized as bright fluorescent dots, with each dot representing one interaction event. As negative control one of the primary antibodies was not added therefore, no positive signals were obtained from that assay). The tissues were visualized using a confocal microscope system (LEICA TCS SP5).

### In situ hybridizations

Protocol was performed according to ^104^. Imaginal discs and brains were dissected and fixed in 4% formaldehyde for 20 min at room temperature, washed in PBS-0.1% Tween (PBT) and re-fixed for 20 min at room temperature with 4% formaldehyde and 0.1% Tween. After three washes in PBT, discs were stored at −20°C in hybridization solution (HS; 50% formamide, 5× SSC, 100 μg/ml salmon sperm DNA, 50 μg/ml heparin and 0.1% Tween). Disc were pre-hybridized for 2 h at 55°C in HS and hybridized with digoxigenin-labelled RNA probes at 55°C. The probes were previously denaturalized at 80°C for 10 min. After hybridization, discs were washed in HS and PBT, and incubated for 2 h at room temperature in a 1:4000 dilution of anti-DIG antibody (Roche). After incubation, the discs were washed in PBT and the detection of probes was carried out using NBT and BCIP solution (Roche). The discs were mounted in 70% glycerol. Images were acquired with a Leica DM750 microscope and Leica MC170HD camera and LASv4.8 software. The probes were generated from the cDNAs RE026007 *(wg)* and LD32066 *(fz1)* from the Expression Sequence Tags (EST) collection of the Berkeley Drosophila Genome Project.

### TEM

Transmission electron microscopy (TEM) was performed in CNS of 3rd instar larvae with horseradish peroxidase (HRP) genetically driven to glial cells. Brains were fixed in 4% formaldehyde in PBS for 30 min at room temperature, and washed in PBS, followed by an amplification of HRP signal using the ABC kit (Vector Laboratories) at room temperature. After developing with DAB brains were washed with PBS and fixed with 2% glutaraldehyde, 4% formaldehyde in PBS for 2h at room temperature. After washing in phosphate buffer the samples were postfixed with OsO4 1% in 0.1 M phosphate buffer, 1% K3[Fe(CN)6] 1h at 4°C. After washing in dH2O, Brains were incubated with tannic acid in PBS for 1min at room temperature then washed in PBS for 5min and dH2O 2×5min. Then the samples were stained with 2% uranyl acetate in H2O for 1h at room temperature in darkness followed by 3 washes in H2O2d. Brains dehydrated in ethanol series (30%, 50%, 70%, 95%, 3×100% 10 min each at 4°C). Infiltration: samples were incubated in EtOH: propylene’s OXID (1:1;V.V) for 5 min, propylene’s OXID 2×10min, propylene’s OXID:Epon (1:1) for 45 min, Epon 100% in agitation for 1 h and Epon 100% in agitation overnight. Then change to Epon 100% for 2-3 h. Finally encapsulate the samples in BEEM capsules and polymerize 48h at 60°C^105^.

### Imaging

Fluorescent labeled samples were mounted in Vectashield mounting media with DAPI (Vector Laboratories) and analyzed by Confocal microscopy (LEICA TCS SP5/SP8). Images were processed using Leica LAS AF Lite and Fiji (Image J 1.50e). Images were assembled using Adobe Photoshop CS5.1.

### Quantifications and Statistical Analysis

Relative Wg, Fz1, Arm, WNT1 and βCatenin staining within brains was determined from images taken at the same confocal settings. Average pixel intensity was measured using measurement log tool from Fiji 1.51g and Adobe Photoshop CS5.1. Average pixel intensity was measured in the glial tissue and in the adjacent neuronal tissue (N=~10 for each sample) and expressed as a ratio. Total average pixel intensity of WNT1 and βCatenin staining within mice brains was measured in the glioma (N=6) and control samples (N=6), to quantify this, single sections were taken from similar z-positions in both control and glioma samples. Glial network volume was quantified using Imaris surface tool (Imaris 6.3.1 software). The number of Proximity ligation assay puncta, HA^+^ puncta, Repo^+^ cells and the number of synaptic active sites was quantified by using the Imaris 6.3.1 software.

The Western Blot bands were quantified by using the Image Studio Lite 5.2 software. Data was analyzed and plotted using GraphPad Prism v7.0.0. A D’Agostino & Pearson normality test was performed and the data found to have a normal distribution were analyzed by a two-tailed t-test with Welch-correction. In the case of multiple comparisons a One-way ANOVA with bonferroni post test was used. The data that did not pass the normality test were submitted to a two-tailed Mann-Whitney U-test or in the case of multiple comparisons a Kruskal-Wallis test with Dunns post test. Error bars represent Standard Error of the Mean, significance was ***p≤0.0001, ** p≤0.001 * p≤0.01, ns=non-significant.

### Viability assays

Flies were crossed and progeny was raised at 25°C under standard conditions. The number of adult flies emerged from the pupae were counted for each genotype. The number of control flies was considered 100% viability and all genotypes are represented relative to controls. Experiments were performed in triplicates.

### Survival assay

Males *Tub-Gal80; Repo-Gal4* were crossed with males bearing a control construct (*UAS–LacZ*) or glioma (*UAS–PI3K^dp110^; UAS-EGFR^λ^*) and raised at 17°C. Progeny bearing a glioma (experimental) or LacZ (control) chromosomes were put at 29°C and viability was calculated as the percentage of surviving flies with respect to the starting number of flies as follows: viability = observed (n° of flies)/starting n° of flies × 100. Six independent vials for glioma (*n*= 6) and control (*n*= 6) were analyzed, with each vial with 10 flies.

### qRT-PCRs

Total RNA was isolated from larvae brains (Trizol, Invitrogen) and cDNAs were synthesized with M-MLV RT (Invitrogen). The following specific probes from applied Biosystems were used: Wingless Dm01814379_m1 and Frizzled1 Dm01793718_g1, RpL32 Dm02151827_g1 was used as housekeeping.

qRT-PCR was performed using Taqman Gene Expression (Applied Biosystems) using a 7500 Real Time PCR System (Applied Biosystems) with cycling conditions of 95°C for 10 min and 40 cycles of 95°C for 15 s and 55°C for 1 min. Each experimental point was performed with samples from two independent crosses and three replicates per experimental point, and differences were assessed with a 2-tailed Student *t* test. Results were normalized using the housekeeping RpL32 and the ΔΔ cycle threshold method and are expressed as the relative change (-fold) of the stimulated group over the control group, which was used as a calibrator. qRT-PCR results were analyzed with 7500 v2.0.6 software (Applied Biosystems).

**Supplementary Figures and VideosFigure S1 (Related to Figure 1): TMs enwrap neurons in GB and cytoneme markers co-localize with glioma network**

(A-B) Control and glioma brains from 3rd instar larvae. Glia is labeled with *UAS-Ihog-RFP* driven by *repo-Gal4* to visualize TMs in glial cells as part of an interconnecting network (red). Glial network is marked with lifeActin-GFP reporter (green) and nuclei are marked with DAPI (blue). Imaris 3D reconstructions are shown in A”’-B”’).

(C-G) Glial network is marked with several cytoneme markers: (C) lifeact-GFP reporter (green and glial nuclei are marked with Repo, magenta), (D) GMA-GFP (green), (E) GPI-YFP (green), (F) GFP-MLC (green), (G) sqh-GFP (green) in a glioma brain. (H-I) Downregulation of neuroglian (*nrg-RNAi*) in glioma brains results in defective TMs.

(J-M) Higher magnifications of control brains (J) showing the glial cytonemes (red) compared with the glioma brains where the TMs overgrow and enwrap neuronal clusters (K). Upon *igl/Gap43* downregulation the glial network does not overgrow or enwrap neuronal clusters (L) and shows a pattern and size similar to the control. Nuclei are marked with DAPI. Arrows indicate glial cytonemes/TMs. (M) A viability assay shows that the lethality induced by the glioma is fully rescued upon knockdown of *Gap43/igl*. Nuclei are marked with DAPI. Scale bar size are indicated in this and all figures. Genotypes: **Figure S1**

(A) *w; lifeActin-GFP; repo-Gal4, UAS-ihog-RFP/UAS-lacZ*, (B-C) *UAS-dEGFR^λ^, UAS-dp110^CAAX^; lifeActin-GFP; repo-Gal4, UAS-ihog-RFP*, (D) *UAS-dEGFR*^λ^*, UAS-dp110^CAAX^; UAS-GMA-GFP; repo-Gal4, UAS-ihog-RFP*, (E) *UAS-dEGFR^λ^, UAS-dp110^CAAX^; UAS-GPI-YFP/Gal80^ts^; repo-Gal4, UAS-myrRFP*, (F) *UAS-dEGFR^λ^, UAS-dp110^CAAX^; Gal80^ts^; repo-Gal4, UAS-myrRFP/ UAS-GFP-sls(MLC)*, (G) *UAS-dEGFR^λ^, UAS-dp110^CAAX^; Gal80^ts^; repo-Gal4, UAS-myrRFP/ Sqh-GFP*, (H-I) *UAS-dEGFR*^λ^*, UAS-dp110^CAAX^; UAS-nrg-RNAi; repo-Gal4, UAS-ihog-RFP*, (J) *w; repo-Gal4, UAS-ihog-RFP/UAS-lacZ*, (K) *UAS-dEGFR*^λ^*, UAS-dp110^CAAX^;; repo-Gal4, UAS-ihog-RFP*, (L) *UAS-dEGFR^λ^, UAS-dp110^CAAX^;; repo-Gal4, UAS-ihog-RFP/UAS-igl-RNAi*

**Figure S2 (Related to Figure 1): igl/*Gap43* Knockdown does not show effects in the number of synapses in the NMJ, in the glial network or in the viability of the flies**

(A-B) Glial cells are stained with Repo (green) and the number of glial cells are quantified in the following genotypes: *Control, Glioma* showing an increase in Repo+ cells, *Glioma;lacZ* and *Glioma;yellow-RNAi* showing a similar number of Repo+ cells to Glioma alone. (C-D) Upon *igl* knockdown by *RNAi* in normal brains, the glial network is similar to the control. Glial cells are marked by Repo in green. Nuclei are marked by DAPI. (E-F) Neurons from the larval neuromuscular junction are stained with Nc82 showing the synaptic active sites. Upon knockdown of *igl* the number of synapses marked by Nc82 (green) is similar to the control. (F) Graph showing the quantification of the synapse number. (G) A viability assay shows that the knockdown of *igl* does not alter the % viability of male and female flies. Error bars show S.D. *** P<0.0001 or ns for non-significant. Genotypes: (A) *UAS-dEGFR**^λ^**, UAS-dp110^CAAX^; UAS-yellow-RNAi; repo-Gal4, UAS-ihog-RFP*, (B) 1. *w; repo-Gal4, ihog-RFP/UAS-lacZ* 2. *UAS-dEGFR****^λ^****, UAS-dp110^CAAX^;; repo-Gal4, UAS-ihog-RFP* 3. *UAS-dEGFR****^λ^****, UAS-dp110^CAAX^; UAS-lacZ; repo-Gal4, UAS-ihog-RFP* 4. *UAS-dEGFR****^λ^****, UAS-dp110^CAAX^; UAS-yellow-RNAi; repo-Gal4, UAS-ihog-RFP*, (C-D) *w; repo-Gal4, UAS-ihog-RFP/UAS-igl-RNAi*, (E-F) 1. *w; UAS-CD8-GFP; D42-Gal4/UAS-igl-RNAi* 2. *w; UAS-CD8-GFP; D42-Gal4/UAS-lacZ*, (G) 1. *w; repo-Gal4, UAS-ihog-RFP/UAS-lacZ* 2. *w; repo-Gal4, UAS-ihog-RFP/UAS-igl-RNAi*

**Figure S3 (Related to Figure 2): Cases of human GB patients with mutations in WNT or FZD.**

Complete analysis of mutations in human GB samples from COSMIC database http://cancer.sanger.ac.uk/cosmic (A) and Cancer Genome Atlas (TGAC) for transcriptional targets of WNT pathway (B), WNT ligands (C) and FZD receptors (D), data are represented in percentage out of 902 or 922 samples. The total number of cases with mutations in any WNT or FZD gene is shown in red. Genes from WNT and FZD family without mutations in GBs is shown in the bottom.

**Figure S4 (Related to Figure 2): Fz2 remains normal in glioma brains.**

(A) Wing imaginal control disc stained with Fz2 (green) showing the expected endogenous localization pattern. Brains from 3rd instar larvae displayed at the same scale. Glia is labeled with *UAS-Ihog-RFP* driven by *repo-Gal4* to visualize active filopodia/TMs in glial cells, and stained with Fz2 (green). (B) Fz2 is homogeneously distributed in control sections, (C) Fz2 is homogeneously distributed in glioma brain sections, similar to the control. Nuclei are marked by DAPI (blue). Scale bar size is indicated in the figure. Glial cell bodies and membranes are labeled with *UAS-myrRFP* (red) driven by *repo-Gal4* and stained with GFP antibody to visualize a tagged form of endogenous Fz1 protein. Genotypes: (A) *w; UAS-lacZ*, (B) *w;; repo-Gal4, ihog-RFP/UAS-lacZ*, (C) *UAS-dEGFR^λ^, UAS-dp110^CAAX^;; repo-Gal4, UAS-ihog-RFP*

**Figure S5 (Related to Figure 2 and 7): Validation of tools, RNAis and antibodies**

(A) *UAS-Fz1-RNAi* tool was validated in epithelial tissue, wing imaginal discs. Upon *UAS-Fz1-RNAi* expression in the posterior compartment (marked with GFP), there is a reduction in Fz1 protein signal compared with the anterior compartment from the same tissue. (B) *UAS-Wg-RNAi* tool was validated in epithelial tissue, wing imaginal discs. Upon *UAS-Wg-RNAi* expression in the posterior compartment (marked with GFP), there is a reduction in Wg protein signal compared with the anterior compartment from the same tissue. (C) *lexAop-Fz1* tool was validated in brain tissue. Upon ectopic Fz1 expression in the neurons (driven by *ELAV-LexA* marked with GFP), there is an increase in Fz1 protein signal compared with the rest of the brain tissue. (D) Upon ectopic Fz1 expression in the neurons (driven by *ELAV-LexA>lexAop-Fz1* marked with GFP), there is an increase in active Arm protein signal compared with the rest of the brain tissue.

Genotypes: (A) w; *Gal80^ts^/UAS-Fz1-RNAi; en-Gal4, UAS-GFP*, (B) w; *Gal80^ts^/UAS-wg-RNAi; en-Gal4, UAS-GFP*, (C-E) w; *lexAop-fz1/ elav-lexA, lexAop-CD8-GFP*

**Figure S6 (Related to Figure 3): Wg and Fz1 transcription levels are similar between controls and gliomas**

(A) qPCRs with *RNA* extracted from control and glioma larvae showing no change in the transcription (*mRNA* levels)of *wg* or *fz1*. (B) Western blot of samples extracted from control and glioma larvae showing no change in the amount of Fz1 or Fz2 protein. Error bars show S.D. ns for non-significant. (C) In situ hybridization experiments for Wg and Fz1 in controls and gliomas showing no change in the transcription (*mRNA* levels) of *wg* or *fz1. Genotypes:* (A-C) 1. *w;; repo-Gal4, ihog-RFP/UAS-lacZ* 2. *UAS-dEGFR^λ^, UAS-dp110^CAAX^;; repo-Gal4, UAS-ihog-RFP*

**Figure S7 (Related to Figure 4): Wg signaling pathway is active in glioma cells**

Larval brain sections with glial network labeled in red and stained with Cyt-Arm (green). (A) Knockdown of *Fz1* in glioma brains showing a homogeneous Cyt-Arm distribution similar to the control. Quantification of Cyt-Arm staining ratio between Ihog+ and Ihog^−^ domains is shown in principal Figure 5D. (B-G) Glial cell bodies and membranes are labeled with myrRFP or ihog-RFP (red) driven by *repo-Gal4*. Wg signaling pathway reporters *tsh-lacZ* stained with anti-bGal (B-C), *fz4-GFP* (D-E) and *dally-lacZ* stained with anti-bGal (F-G) show activation of the pathway in glial transformed cells. Genotypes: (A) *UAS-dEGFR^λ^, UAS-dp110^CAAX^; UAS-Fz1-RNAi; repo-Gal4, UAS-ihog-RFP*, (B) *w*;; *repo-Gal4, UAS-myrRFP/tsh-lacZ*, (C) *UAS-dEGFR^λ^, UAS-dp110^CAAX^;; repo-Gal4, UAS-myrRFP/tsh-lacZ*, (D) *w;; repo-Gal4, UAS-myrRFP/ fz4-GFP*, (E) *UAS-dEGFR^λ^, UAS-dp110^CAAX^;; repo-Gal4, UAS-myrRFP/ fz4-GFP*, (F) *w;; repo-Gal4, UAS-myrRFP/ dally-lacZ*, (G) *UAS-dEGFR^λ^, UAS-dp110^CAAX^;; repo-Gal4, UAS-myrRFP/ dally-lacZ*

**Figure S8 (Related to Figure 4): Wg signaling pathway is active in human glioma cells**

(A-D) A series of grade II and III GB images from S24 xenografts brain sections stained with WNT1 and ß-Catenin show an increase of these signals in grade III when compared with grade II brain sections, indicating that the accumulation of WNT1 and ß-Catenin correlates with the progression of the GB, quantified in (E-F) (G-H) Technical immunohistofluorescence negative control in NMRI nude control mice brains stained only with the corresponding secondary antibodies showing the background unspecific signal. Nuclei are marked by DAPI (blue).

**Figure S9 (Related to Figure 2, 4 and 6): Adult *Drosophila* gliomas**

(A) Survival curve of adult control or glioma flies after a number of days of glioma induction and progression. (B-H) Adult brain sections 7 days after glioma induction with Glial cells are labeled with *UAS-myr-RFP* to visualize the glial network and stained with Cyt-Arm, Fz1 and Wg antibodies. (B-C, D) Cyt-Arm staining specifically marks the mushroom bodies and it is homogeneously distributed in the rest of the brain tissue in control sections and accumulates in the neurons cytoplasm where it is inactive in glioma brains. Quantification of Neuron/Glia Cyt-Arm staining ratio between RFP+ and RFP^−^ domains (D). (B’-C’, E) Fz1 staining show homogeneous localization in the control brains (B’) in blue. In the glioma brains Fz1 accumulates in the glial transformed cells (C’), Glia/Neuron Fz1 average pixel intensity ratio quantification is shown in (E). (F-H) Wg is homogeneously distributed in control brains, with a slight accumulation in the RFP+ structures. Wg accumulates in the glioma network similar to the larval brains. Glia/Neuron Wg average pixel intensity ratio quantification is shown in (H). (I) Graph showing synapse number quantification of adult NMJs from control flies and glioma-bearing flies. Error bars show S.D. *** P<0.0001 or ns for non-significant. Genotypes: (B, F) w; *Gal80^ts^; repo-Gal4, UAS-myrRFP/UAS-lacZ*, (C, G) *UAS-dEGFR^λ^, UAS-dp110^CAAX^; Gal80^ts^; repo-Gal4, UAS-myrRFP*

**Figure S10 (Related to Figure 7): Restoration of the glia-neuron Wg/Fz1 signaling equilibrium inhibits glioma progression**

(A-D) Brains from 3rd instar larvae displayed at the same scale. Glia is labeled with *UAS-Ihog-RFP* driven by *repo-Gal4* to visualize active filopodia in glial cells, and stained with Wg or Cyt-Arm (green). Neurons are labeled with *lexAop-CD8-GFP* driven by *elav-lexA* (blue). Fz1 overexpression in neurons (blue) restore homogeneous Wg (grey or green in the merge) (B) and Cyt-Arm (D) protein distribution (green) in the brain, compared to (A, C) where the *elav-lexA* is not present in the glioma brains, Nuclei are marked by DAPI (blue) in (A, C). Genotypes: (A, C) *UAS-dEGFR^λ^, UAS-dp110^CAAX^; lexAop-Fz1; repo-Gal4, UAS-ihog-RFP*, (B, D) *UAS-dEGFR^λ^, UAS-dp110^CAAX^; lexAop-Fz1/elav-lexA, lexAop-CD8-GFP; repo-Gal4, UAS-ihog-RFP*

**Video S1: Control Network**

3D video reconstruction of control brains with glia labeled with *ihog-RFP* (*repo>ihog-RFP*) in red (grey in the 3D reconstruction) to visualize cytoneme structures in glial cells as part of an interconnecting network.

**Video S2: Glioma TMs**

3D video reconstruction of Glioma brains with glia labeled with *ihog-RFP* (*repo>ihog-RFP*) in red (grey in the 3D reconstruction) to visualize TMs structures in glial cells as part of an interconnecting network. In glioma brains, the TMs expand across the brain and form perineuronal nests.

**Video S3: Glioma *igl-RNAi* Network**

3D video reconstruction of Glioma; *igl-RNAi* brains with glia labeled with *ihog-RFP* (*repo>ihog-RFP*) in red (grey in the 3D reconstruction) to visualize TM structures in glial cells as part of an interconnecting network. Upon *igl* downregulation the glial network does not overgrow or enwrap neuronal clusters and shows a pattern and size similar to the control.

**Video S4: Control LifeActin**

3D video reconstruction of control brains from 3rd instar larvae. Glia is labeled with *UAS-Ihog-RFP* driven by *repo-Gal4* to visualize cytonemes in glial cells as part of an interconnecting network (red). Glial network is marked with lifeActin-GFP reporter (green) and nuclei are marked with DAPI (blue).

**Video S5: Glioma LifeActin**

3D video reconstruction of gliomal brains from 3rd instar larvae. Glia is labeled with *UAS-Ihog-RFP* driven by *repo-Gal4* to visualize TMs in glial cells as part of an interconnecting network (red). Glial network is marked with lifeActin-GFP reporter (green) and nuclei are marked with DAPI (blue). Glial TMs enwrap clusters of neurons in individual GB perineuronal nests.

**Video S6: Climbing assay**

Video of *Drosophila* adult negative geotaxis behavior analysis (climbing assay), as an indication for possible motor defects associated with neurodegeneration. The results showed symptoms of neurodegeneration in glioma flies (right tube) compared to controls (left tube).

