## Supplementary figures and images for "WNT vampirization by glioblastoma leads to tumor growth and neurodegeneration"

### Supplementary file 7

Portela_FigS1

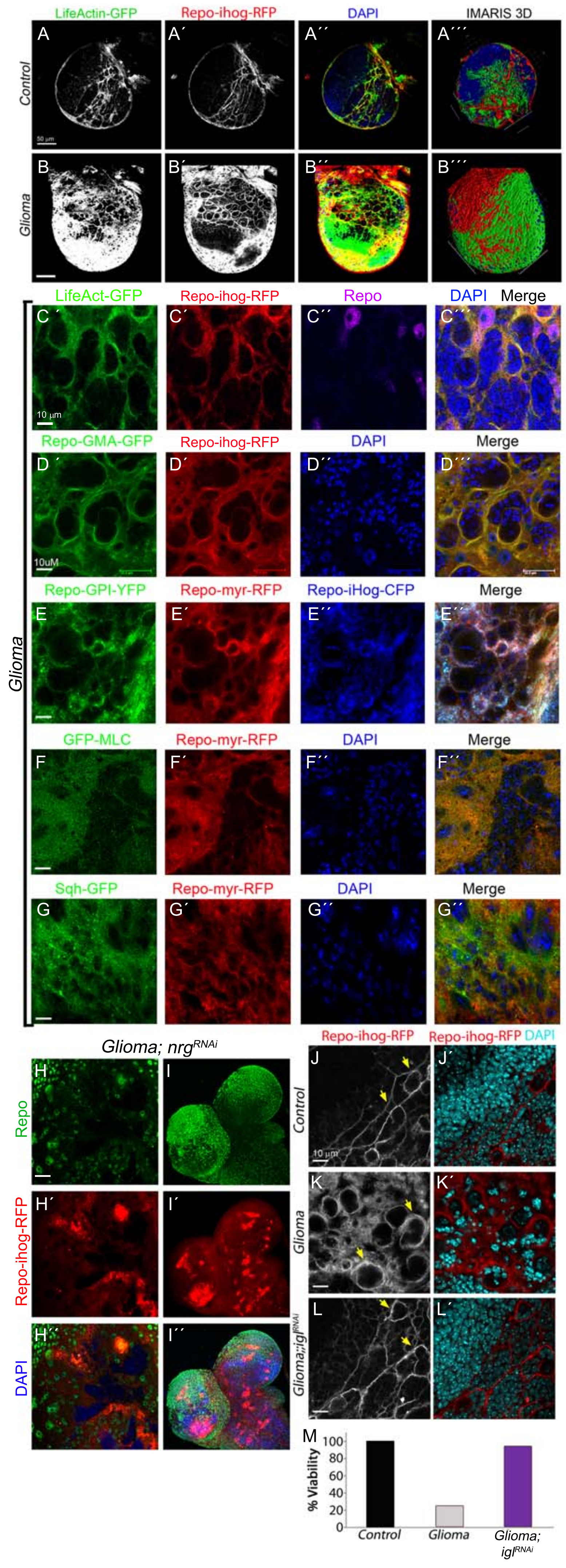

### Supplementary file 8

Portela_FigS2

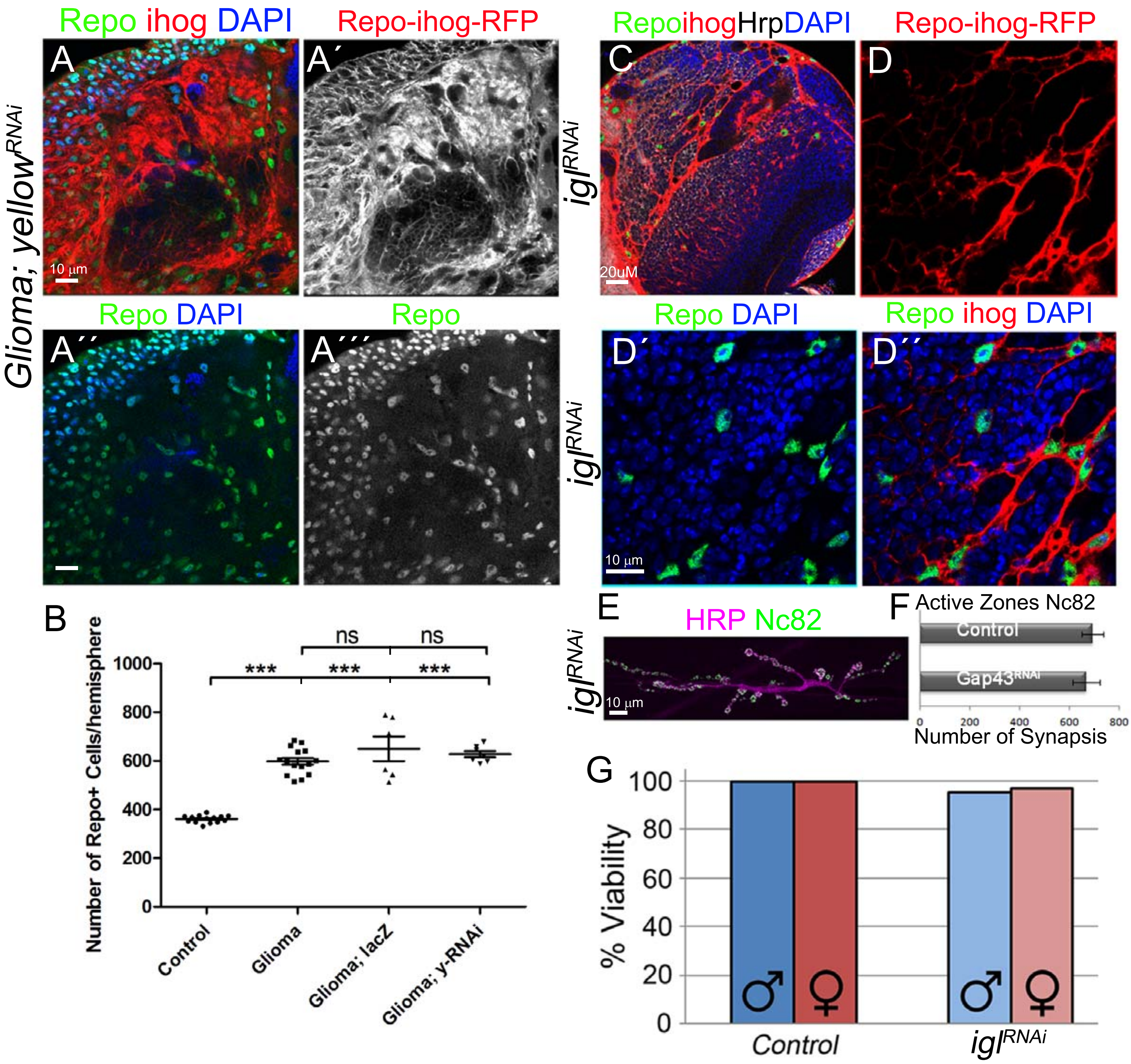

### Supplementary file 10

Portela_FigS4

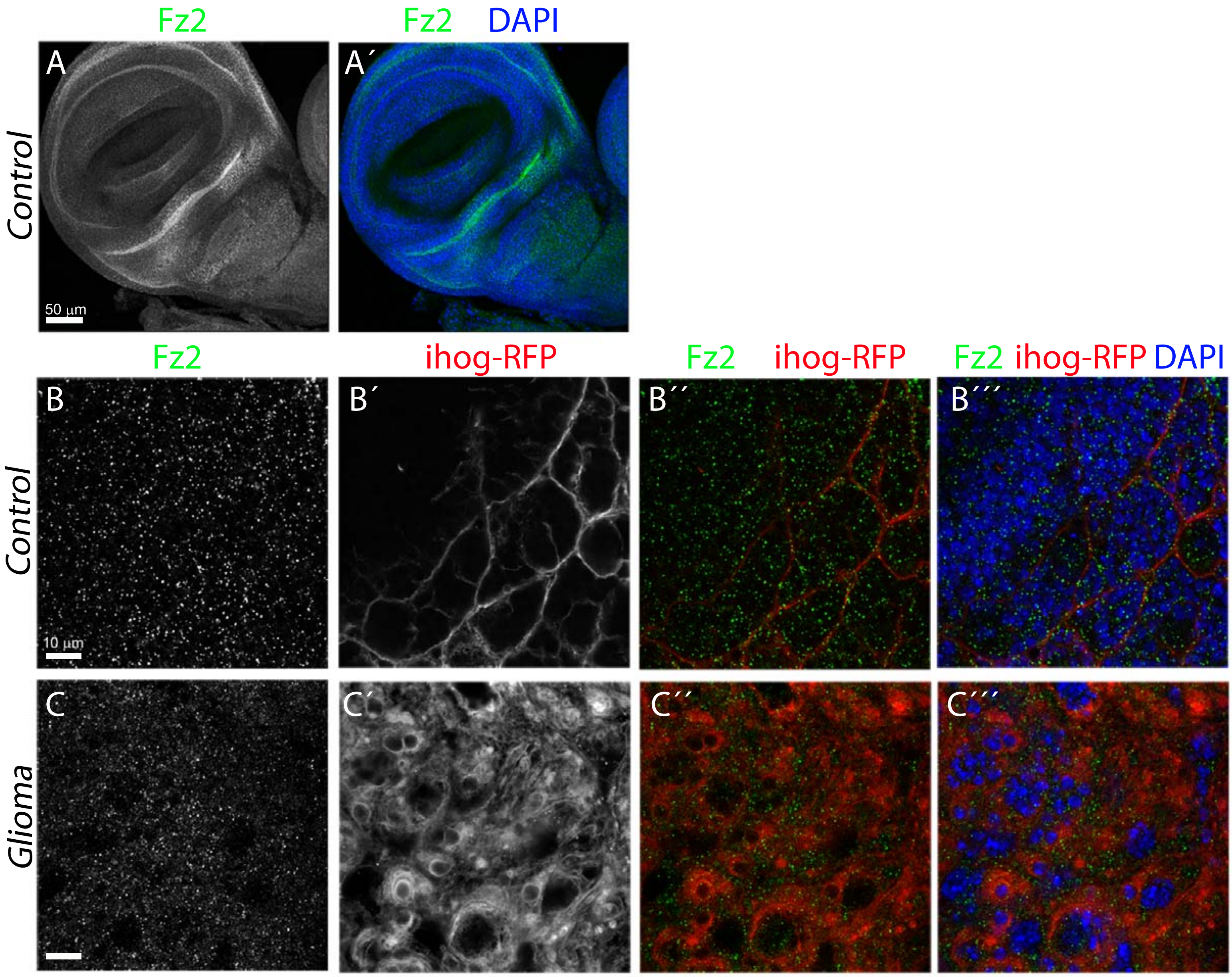

### Supplementary file 11

Portela_FigS5

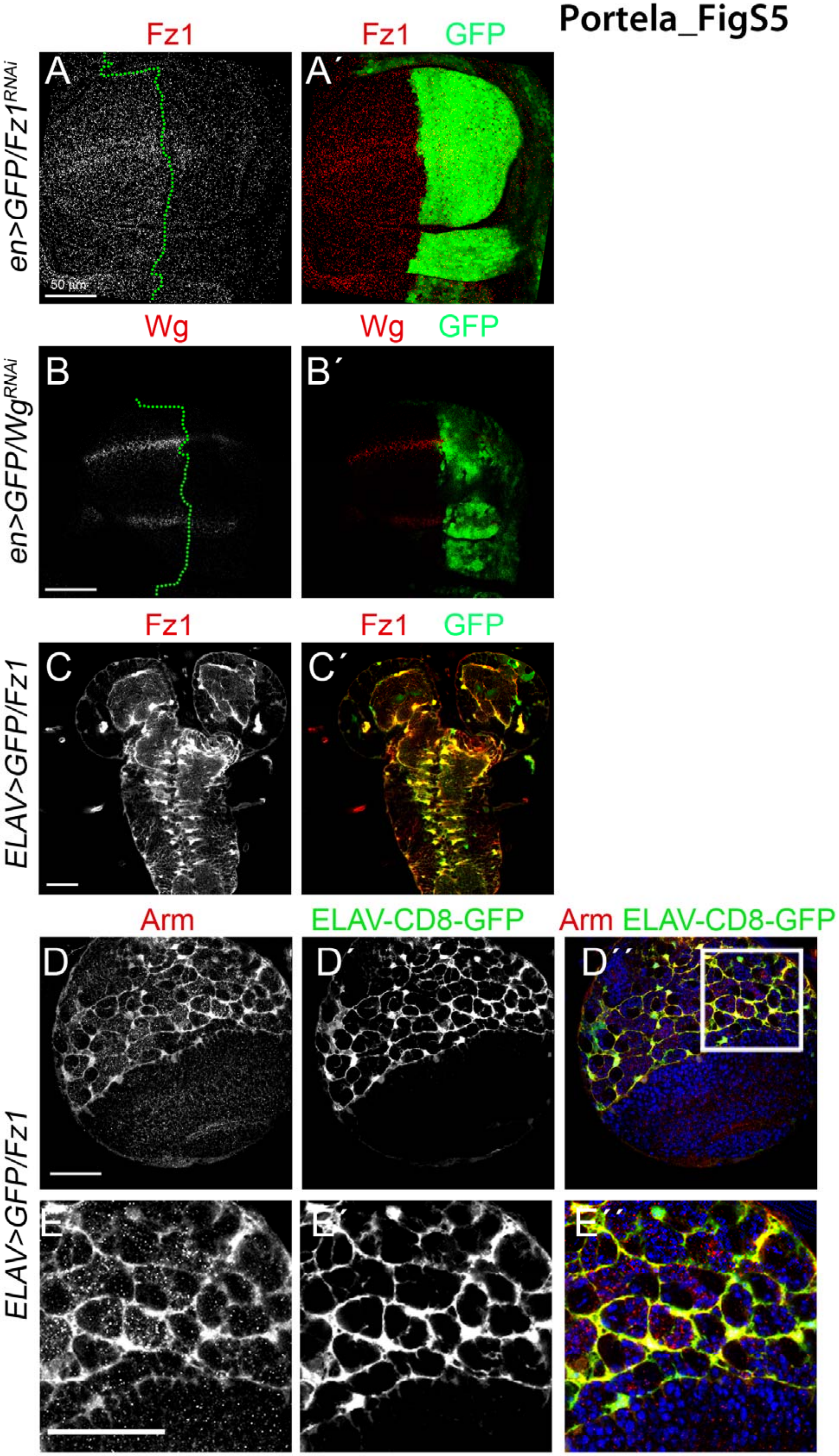

### Supplementary file 12

**Portela_FigS6**

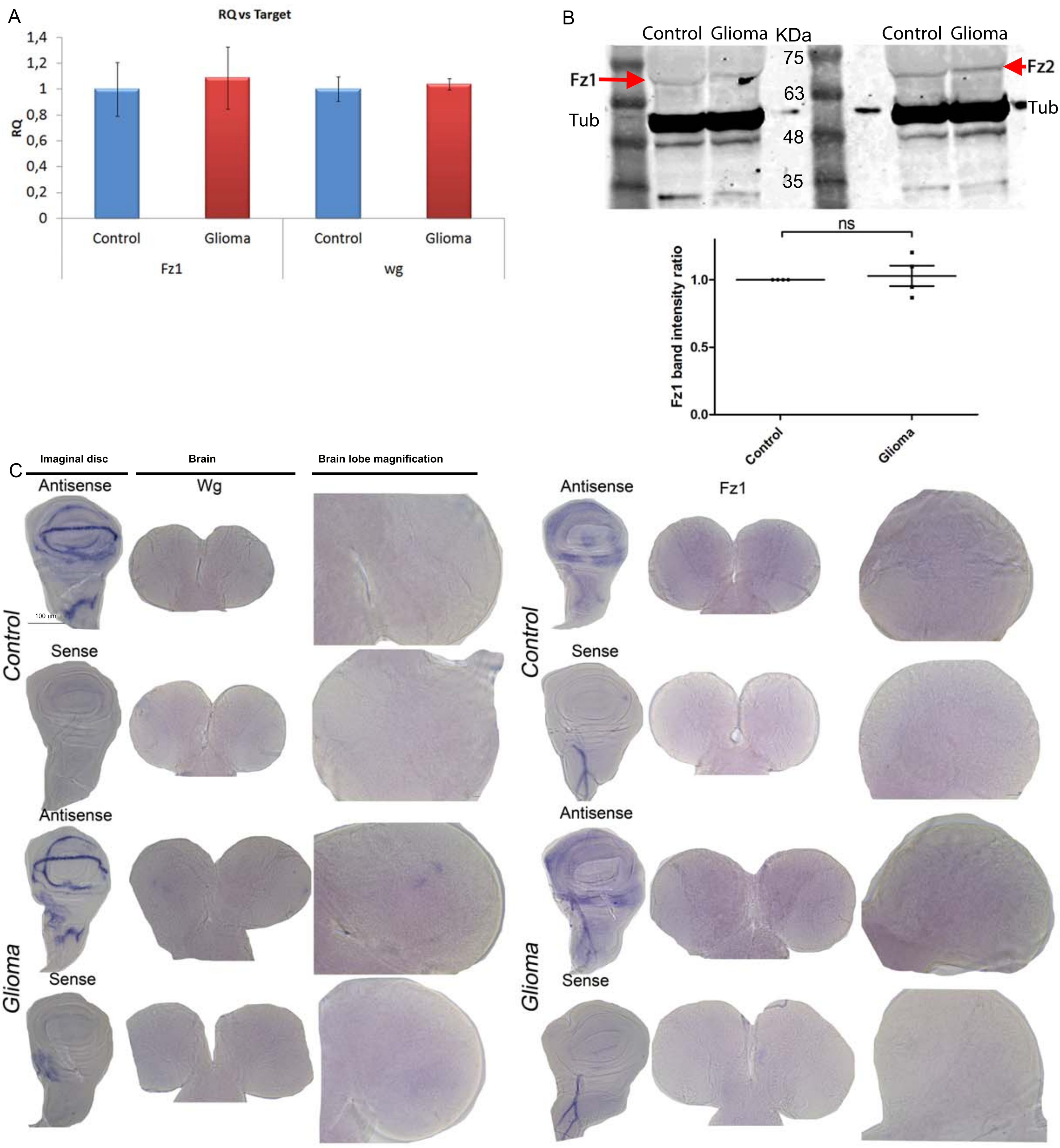

### Supplementary file 13

#### Portela_FigS7

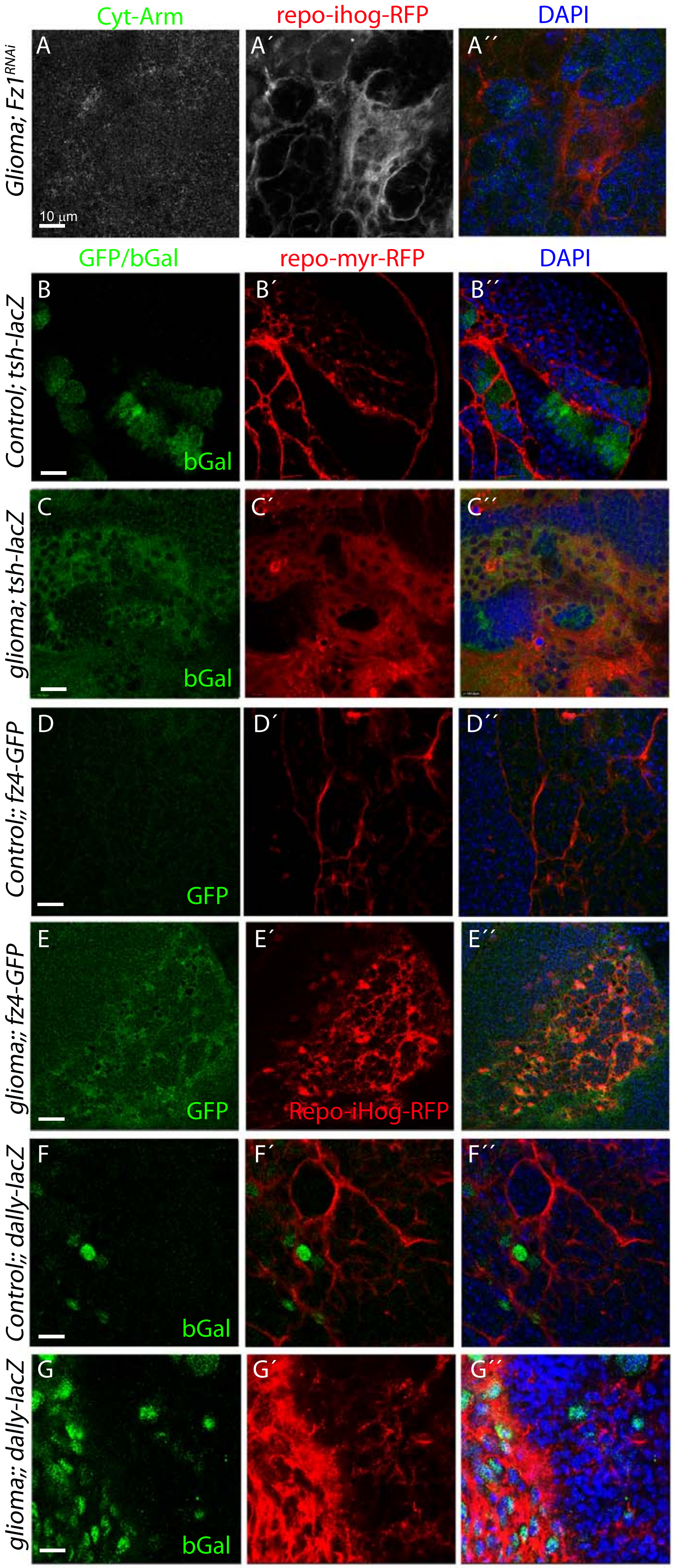

### Supplementary file 14

#### Portela_FigS8

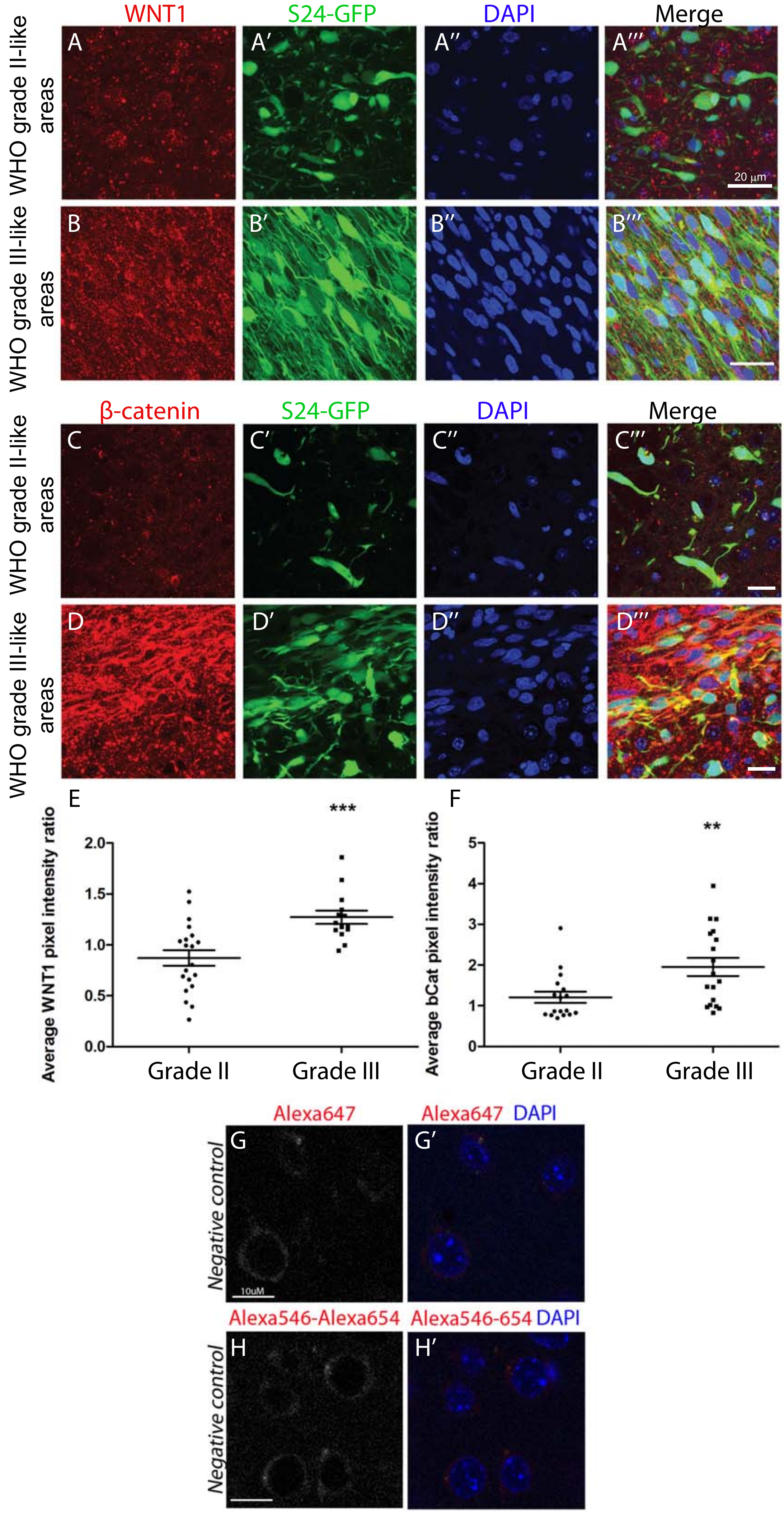

### Supplementary file 15

Portela_FigS9

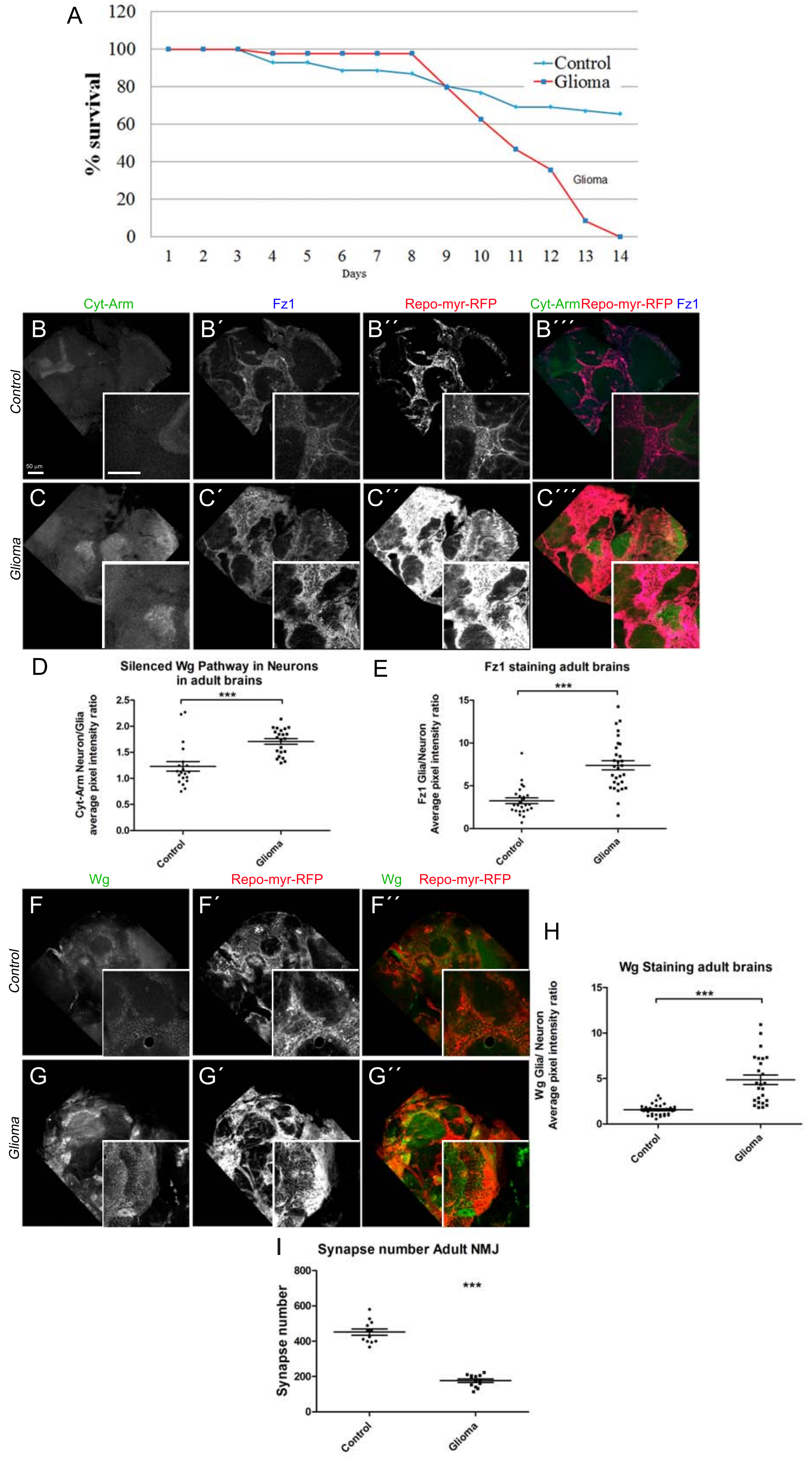

### Supplementary file 16

Portela_FigS10

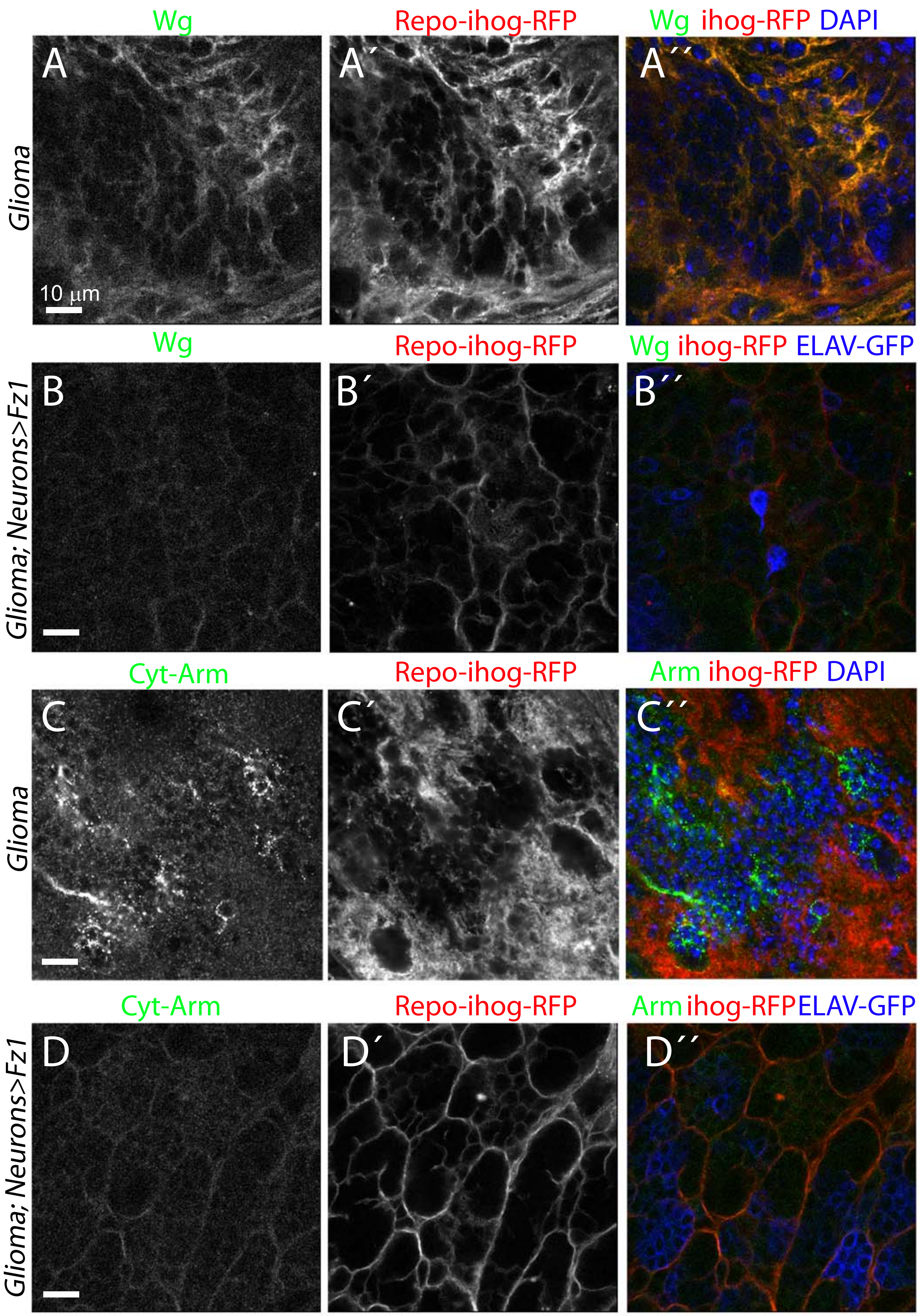
