## Supplementary material for "WNT vampirization by glioblastoma leads to tumor growth and neurodegeneration"

**Portela_FigS3**

# A

### COSMIC Database

| *Gene* | *Cases with mutations* | *Number of analyzed GBMs* | *% of mutantions* |
|---|---|---|---|
| WNT2 | 6 | 922 | 0,650759219 |
| WNT2B | 2 | 922 | 0,21691974 |
| WNT2B_ENST00000369686 | 1 | 902 | 0,110864745 |
| WNT3 | 1 | 922 | 0,10845987 |
| WNT5B_ENST00000397196 | 1 | 902 | 0,110864745 |
| WNT8A | 1 | 922 | 0,10845987 |
| WNT8B_ENST00000343737 | 1 | 902 | 0,110864745 |
| WNT9A | 3 | 922 | 0,32537961 |
| WNT9B | 2 | 922 | 0,21691974 |
| WNT9B_ENST00000575372 | 2 | 902 | 0,22172949 |
| WNT10A | 2 | 922 | 0,21691974 |
| WNT10B | 1 | 922 | 0,10845987 |
| WNT11 | 1 | 922 | 0,10845987 |
| SUM WNT | | | 2,615061252 |
| FZD1 | 3 | 922 | 0,32537961 |
| FZD3 | 3 | 922 | 0,32537961 |
| FZD5 | 1 | 922 | 0,10845987 |
| FZD6 | 3 | 923 | 0,325027086 |
| FZD7 | 1 | 922 | 0,10845987 |
| FZD8 | 3 | 902 | 0,332594235 |
| FZD9 | 4 | 922 | 0,433839479 |
| FZD10 | 5 | 922 | 0,542299349 |
| SUM FZD | | | 2,501439108 |
| ASCL1 | 1 | 922 | 0,10845987 |
| DVL2 | 2 | 922 | 0,21691974 |
| FOXM1 | 5 | 922 | 0,542299349 |
| **Genes without mutations in GBMs** | | | |
| WNT1 | 0 | 922 | |
| WNT3A | 0 | 922 | |
| WNT3A_ENST00000366753 | 0 | 902 | |
| WNT4 | 0 | 922 | |
| WNT4_ENST00000374655 | 0 | 902 | |
| WNT5A | 0 | 922 | |
| WNT5B | 0 | 922 | |
| WNT6 | 0 | 922 | |
| WNT6_ENST00000233948 | 0 | 902 | |
| WNT7A | 0 | 922 | |
| WNT7B | 0 | 922 | |
| WNT8B | 0 | 922 | |
| WNT16 | 0 | 922 | |
| WNT16_ENST00000361301 | 0 | 902 | |
| FZD2 | 0 | 922 | |
| FZD4 | 0 | 922 | |

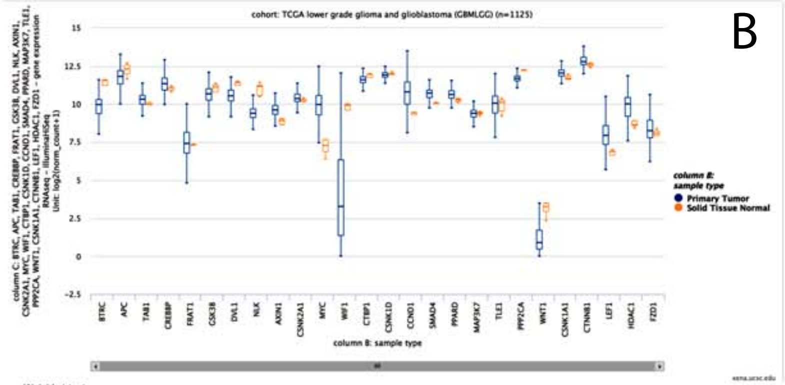


Welch's t-test

| | | | | | | |
|---|---|---|---|---|---|---|
| BTRC<br>t = -4.799<br>p = 0.008313 | APC<br>t = -2.937<br>p = 0.04003 | TAB1<br>t = 0.8518<br>p = 0.4414 | CREBBP<br>t = 4.042<br>p = 0.009591 | FRAT1<br>t = 1.549<br>p = 0.1892 | GSK3B<br>t = -1.270<br>p = 0.2716 | DVL1<br>t = -15.32<br>p = 0.00001040 |
| NLK<br>t = -8.926<br>p = 0.0007450 | AXIN1<br>t = 2.989<br>p = 0.03962 | CSNK2A1<br>t = 2.677<br>p = 0.04488 | MYC<br>t = 6.296<br>p = 0.003012 | WIF1<br>t = -5.076<br>p = 0.006603 | CTBP1<br>t = -5.677<br>p = 0.003065 | CSNK1D<br>t = -0.7610<br>p = 0.4874 |
| CCND1<br>t = 4.926<br>p = 0.006881 | SMAD4<br>t = 8.597<br>p = 0.0005964 | PPARD<br>t = 7.193<br>p = 0.0009659 | MAP3K7<br>t = -0.6891<br>p = 0.5226 | TLE1<br>t = 0.3668<br>p = 0.7315 | PPP2CA<br>t = -6.207<br>p = 0.003009 | |
| WNT1<br>t = -4.209<br>p = 0.01336 | CSNK1A1<br>t = 5.543<br>p = 0.004005 | CTNNB1<br>t = 4.675<br>p = 0.006656 | LEF1<br>t = 7.816<br>p = 0.001002 | HDAC1<br>t = 4.405<br>p = 0.01068 | FZD1<br>t = 2.547<br>p = 0.05957 | |

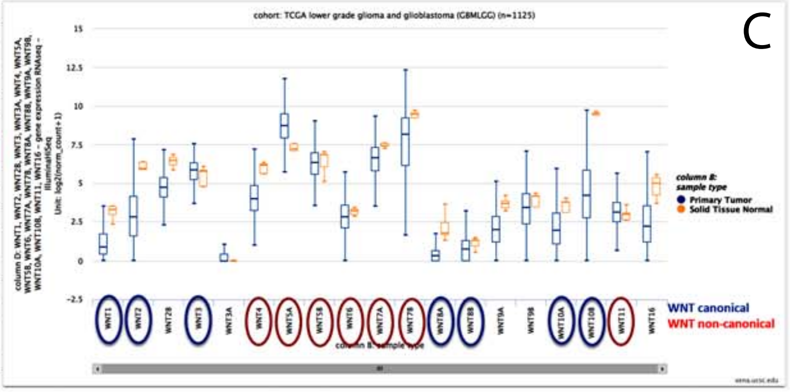


Welch's t-test

| | | | | | | | | | |
|---|---|---|---|---|---|---|---|---|---|
| WNT1<br>t = -4.209<br>p = 0.01336 | WNT2<br>t = -8.086<br>p = 0.0009206 | WNT2B<br>t = -10.12<br>p = 0.0002710 | WNT3<br>t = 1.105<br>p = 0.3291 | WNT3A<br>t = 13.50<br>p = 0.000 | WNT4<br>t = -11.94<br>p = 0.00006337 | WNT5A<br>t = 6.866<br>p = 0.001762 | WNT5B<br>t = -0.3102<br>p = 0.7715 | WNT6<br>t = -2.360<br>p = 0.05761 | |
| WNT7A<br>t = -4.725<br>p = 0.007457 | WNT7B<br>t = -3.700<br>p = 0.01734 | WNT8A<br>t = -4.806<br>p = 0.008486 | WNT8B<br>t = -1.705<br>p = 0.1587 | WNT9A<br>t = -9.615<br>p = 0.0002432 | WNT9B<br>t = -0.1077<br>p = 0.9193 | WNT10A<br>t = -2.320<br>p = 0.07898 | WNT10B<br>t = -13.69<br>p = 0.00008213 | WNT11<br>t = 1.062<br>p = 0.3412 | WNT16<br>t = -6.564<br>p = 0.001849 |

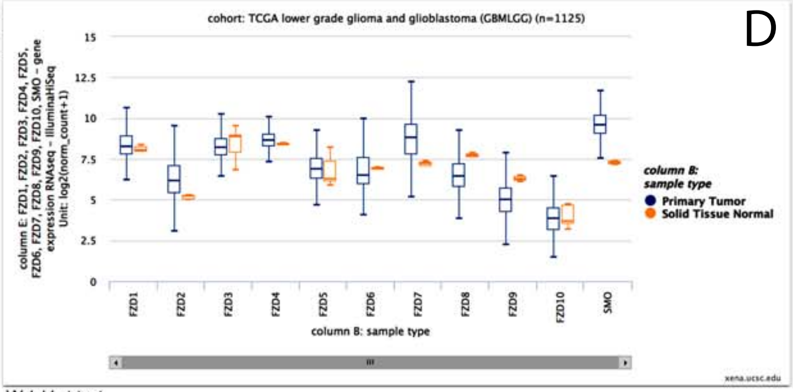


Welch's t-test

| | | | | | |
|---|---|---|---|---|---|
| FZD1<br>t = 2.547<br>p = 0.05957 | FZD2<br>t = 7.873<br>p = 0.0003746 | FZD3<br>t = -0.4067<br>p = 0.7048 | FZD4<br>t = 1.944<br>p = 0.1212 | FZD5<br>t = 0.5511<br>p = 0.6105 | |
| FZD6<br>t = 0.2234<br>p = 0.8329 | FZD7<br>t = 11.55<br>p = 0.00001613 | FZD8<br>t = -11.42<br>p = 0.00001797 | FZD9<br>t = -16.90<br>p = 2.315e-7 | FZD10<br>t = -0.3254<br>p = 0.7606 | SMO<br>t = 19.65<br>p = 0.000007692 |
